# Unveiling the Genetic Tapestry: Rare Disease Genomics of Spinal Muscular Atrophy and Phenylketonuria Proteins

**DOI:** 10.1101/2023.11.27.568432

**Authors:** Debaleena Nawn, Sk. Sarif Hassan, Elrashdy M. Redwan, Tanishta Bhattacharya, Pallab Basu, Kenneth Lundstrom, Vladimir N. Uversky

**Affiliations:** Indian Research Institute for Integrated Medicine (IRIIM), Unsani, Howrah, 711302, West Bengal, India; Department of Mathematics, Pingla Thana Mahavidyalaya, Maligram, Paschim Medinipur, West Bengal, India; Biological Science Department, Faculty of Science, King Abdulaziz University, Jeddah, Saudi Arabia, Therapeutic and Protective Proteins Laboratory, Protein Research Department, Genetic Engineering and Biotechnology Research Institute, City of Scientific Research and Technological Applications, New Borg EL-Arab, 21934, Alexandria, Egypt; Developmental Genetics (Dept III), Max Planck Institute for Heart and Lung Research, Ludwigstrabe 43, 61231, Bad Nauheim, Germany; School of Physics, University of the Witwatersrand, Johannesburg, Braamfontein 2000, South Africa; Adjunct Faculty, Woxsen School of Sciences, Woxsen University, Hyderabad – 500 033, Telangana, India; PanTherapeutics, Rte de Lavaux 49, CH1095 Lutry, Switzerland; Department of Molecular Medicine, Morsani College of Medicine, University of South Florida, Tampa, FL 33612, USA

**Keywords:** Phenylalanine 4-monooxygenase (PAH) and Survival motor neuron (SMN1), SMA, PKU, Agglomerated Proximal Sequences, Quantitative Genomics, Rare Disease

## Abstract

Rare diseases, defined by their low prevalence, present significant challenges, including delayed detection, expensive treatments, and limited research. This study delves into the genetic basis of two noteworthy rare diseases in Saudi Arabia: Phenylketonuria (PKU) and Spinal Muscular Atrophy (SMA). PKU, resulting from mutations in the phenylalanine hydroxylase (PAH) gene, exhibits geographical variability and impacts intellectual abilities. SMA, characterized by motor neuron loss, is linked to mutations in the survival of motor neuron 1 (SMN1) gene. Recognizing the importance of unveiling signature genomics in rare diseases, we conducted a quantitative study on PAH and SMN1 proteins of multiple organisms by employing various quantitative techniques to assess genetic variations. The derived signature-genomics contributes to a deeper understanding of these critical genes, paving the way for enhanced diagnostics and treatments for disorders associated with PAH and SMN1.

## 1. Introduction

The prevalent definitions of rare diseases revolve around the criteria employed by pharmaceutical regulatory bodies to facilitate the development of therapies for rare conditions, commonly referred to as orphan drugs [1, 2]. These definitions exhibit regional variations. For instance, in the United States, a rare disease is delineated by the 1983 Orphan Drug Act as a condition impacting fewer than 200,000 individuals. As per the National Institutes of Health (NIH), around 7,000 rare diseases impact an estimated 25 to 30 million Americans, translating to one in every ten Americans. The equivalent legislation implemented in the European Union in 2000 designates a disease as rare, when it afflicts less than 1 in 2,000 people [3]. Furthermore, some diseases are ‘rare’ in some demographics or regions, but not in others [4, 5, 6, 7]. Notably, over 80% of these rare diseases have a discernible genetic basis, and roughly 75% of these conditions impact children, according to the Saudi Ministry of Health [8]. Recent statistics from 2023 report 7,362 cases mentioned in the OMIM and Orphanet databases, highlighting the substantial burden of rare diseases [9]. Saudi Arabia alone grapples with an estimated 6,000-8,000 distinct rare diseases, a higher prevalence likely stemming from an increased occurrence of pathogenic alleles linked to consanguineous marriages [8]. This elevated number of rare diseases poses significant challenges, including delayed detection and diagnosis, costly treatment options, and a dearth of pertinent research and scientific studies [10].

Recognizing the growing global importance of understanding and diagnosing rare diseases, various initiatives have been established [11]. For instance, the Orphanet database offers comprehensive information about rare diseases, encompassing their epidemiological distribution, causative factors, and currently available treatment strategies. Additionally, scientific symposiums dedicated to rare diseases have been organized to foster knowledge exchange [12]. Recent years have witnessed pivotal research in smaller cohorts and patient-based studies, shedding light on common mutations underlying various rare diseases, particularly those rooted in genetics [13]. Furthermore, these studies have underscored the role of genetic variations, environmental factors, and geographic disparities in survival in shaping the occurrence and prevalence of rare diseases [14].

Among the rare diseases prevalent in Saudi Arabia, two noteworthy examples are Phenylketonuria (PKU) and Spinal Muscular Atrophies (SMAs). PKU is an autosomal recessive genetic disorder resulting from mutations in the phenylalanine hydroxylase (PAH) gene [15]. The PAH gene plays a vital role in converting Phe to Tyr, and mutations in this gene can disrupt this conversion process, leading to the accumulation of Phe in the bloodstream and the formation of phenyl ketone bodies, typically excreted in urine [16, 17, 18, 19].

PKU exhibits geographical variability in its prevalence, affecting approximately 450,000 individuals worldwide [20]. Italy, with a prevalence of 1 in 4,000, and Ireland, with a prevalence of 1 in 4,545, exhibited higher rates than Iran and Jordan, both at 1 in 5,000, as well as Turkey at 1 in 6,667. In contrast, Saudi Arabia (1 in 14,245), Iraq (1 in 14,286), and the United Arab Emirates (1 in 14,493) had lower prevalence rates of PKU [20]. Individuals with PKU are characterized by lower intellectual abilities, and during the neonatal stage, it can lead to developmental delays, epilepsy, altered psychological behavior, tremors, eczema, and an unusual odor [21, 22]. PKU can be categorized into different stages based on the concentration of Phe in the blood, including mild PKU, moderate PKU, and classic PKU, which represents the severe phenotype [17, 23]. Past studies have revealed that the coding sequence of the PAH gene consists of 452 amino acids, spans 1356 base pairs, and has a molecular weight of approximately 52KDa [22]. The PAH gene comprises three distinct domains [24]:

- The Regulatory N-terminal domain (amino acids 1-142)
- The Catalytic central domain (amino acids 143-410)
- The Oligomerization domain, C-terminal, which includes dimerization and tetramerization motifs (amino acids 411-452).

Spinal Muscular Atrophy (SMA) is also an autosomal recessive disorder characterized by the loss of motor neurons in the spinal cord and lower brainstem, leading to muscle weakening and atrophy [25, 26, 27]. Recent research has indicated a significant prevalence of SMA in Saudi Arabia, where the numbers hover around 32 in 100,000 births, exceeding global rates [28, 29]. SMA, like many rare diseases, has a genetic basis [30]. Mutations in the survival of motor neuron 1 (SMN1) gene, particularly the deletion of exon 7, can trigger the development of this condition [31, 32]. SMN1 plays a crucial role in RNA processing and transportation [33]. Depending on the age of onset and the level of motor function that can be achieved, SMA can be categorized into four phenotypes: SMA I, SMA II, SMA III, and SMA IV, with decreasing severity [34]. SMA I accounts for nearly 60% of all SMA cases and can be diagnosed through genetic testing of SMN1 and copy number testing of SMN2 [35]. The clinical severity of Spinal Muscular Atrophy (SMA) is inversely correlated with the copy number of the SMN2 gene. It ranges from severe weakness and paraplegia in infancy to a milder proximal weakness observed in adulthood [36]. In some instances, specific sequence variations within SMN2, such as the c.859G>C substitution in exon 7, have been found. This substitution creates a new exonic splicing enhancer element, leading to an increase in full-length transcripts and resulting in less severe SMA phenotypes. Therefore, SMN2, while generally known for its dose-dependent influence, can also be subject to sequence variations that modify the disease’s severity [37].

Quantitative genomics is pivotal for advancing the diagnosis and understanding of rare diseases, such as PKU and SMA, which have a genetic basis [38, 39]. Utilizing advanced technologies like next-generation sequencing, genomic analysis enables the identification of specific genetic variations responsible for these conditions by pinpointing mutations or alterations in genes [40]. This precision in genetic diagnosis not only enhances accuracy but also paves the way for personalized and targeted treatment strategies [41]. Additionally, genomics significantly contributes to early detection and intervention in rare diseases, allowing clinicians to diagnose conditions before symptoms appear [42]. Early diagnosis is crucial for timely interventions, potentially altering disease trajectories and improving outcomes [43]. Furthermore, genomic data from rare disease patients contribute to ongoing research, fostering collaborations and expediting the development of novel therapies tailored to specific genetic aberrations [44]. In essence, genomics is a cornerstone in the comprehensive understanding, diagnosis, and treatment of rare diseases, offering a route to more precise and personalized medical care [45]. Quantitative genomics of PAH contributes to a more accurate assessment of enzyme activity levels and the severity of PKU [46]. For SMN1, quantitative genomics aids in assessing the copy number variations of the SMN1 gene, crucial for determining the severity of SMA [47, 48].

In this present study, we focused on deriving basic signature-genomics of SMN1 and PAH genes collected from various organisms to gain valuable insights into their fundamental characteristics and variations. The derived signature genomics not only contributes to our knowledge of these critical genes, but also sets the stage for future investigations into targeted therapies, diagnostics, and personalized medicine strategies.

## 2. Data acquisition

For both the proteins Phenylalanine 4-monooxygenase (PAH) and Survival motor neuron (SMN1), BLAST was made using Human PAH (P00439) and Human SMN1 (Q16637), 62 PAH and 50 SMN1 sequences were obtained with at least 95% similarity with Human PAH (P00439) and Human SMN1 (Q16637), respectively. A list of identical (100% similarity) sequences is given in Table 1. Among them, 51 PAH and 45 SMN1 unique proteins were obtained from the UniProt database as listed in Table 2.

**Table 1:** List of identical sequences of PAH and SMN1 proteins from various organisms.

| *Serial No.* | *List of 100% identical sequences* |  |  |  | *Remarks* |
| --- | --- | --- | --- | --- | --- |
| 1 | tr\|A0A0D9S1L9\|PAH_CHLSB | tr\|A0A2K6BHX1\|PAH_MACNE | tr\|A0A2K5NEI1\|PAH_CERAT | tr\|A0A2K5ZL20\|PAH_MANLE | 100% Identical |
| 2 | tr\|A0A2I3S5K6\|PAH_PANTR | tr\|A0A2R8ZJB8\|PAH_PANPA) |  |  | 100% Identical |
| 3 | tr\|A0A2I3SWI0\|PAH_PANTR | tr\|A0A2R8ZCJ2\|PAH_PANPA | tr\|G3QF53\|PAH_GORGO |  | 100% Identical |
| 4 | tr\|A0A2K5HWD5\|PAH_COLAP | tr\|A0A8C9GZW7\|PAH_9PRIM |  |  | 100% Identical |
| 5 | tr\|A0A2K5NDZ8\|PAH_CERAT | tr\|A0A2K5W2Q3\|PAH_MACFA | tr\|A0A2K5ZKY3\|PAH_MANLE |  | 100% Identical |
| 6 | tr\|A0A2R8ZCI5\|PAH_PANPA | tr\|H2Q6R0\|PAH_PANTR |  |  | 100% Identical |
| 7 | tr\|A0A8C4MZ99\|PAH_EQUAS | tr\|F7BKF9\|PAH_HORSE |  |  | 100% Identical |
| 1 | tr\|A0A2I3MMK8\|SMN_PAPAN | tr\|A0A2K6CHD7\|SMN_MACNE | tr\|A0A2K5UX29\|SMN_MACFA |  | 100% Identical |
| 2 | tr\|A0A2K6LS96\|SMN_RHIBE | tr\|A0A2K6QRN8\|SMN_RHIRO |  |  | 100% Identical |
| 3 | tr\|A0A8I5R6V5\|SMN_PAPAN | sp\|Q4R4F8\|SMN_MACFA | tr\|F6V985\|SMN_MACMU |  | 100% Identical |

**Table 2:**
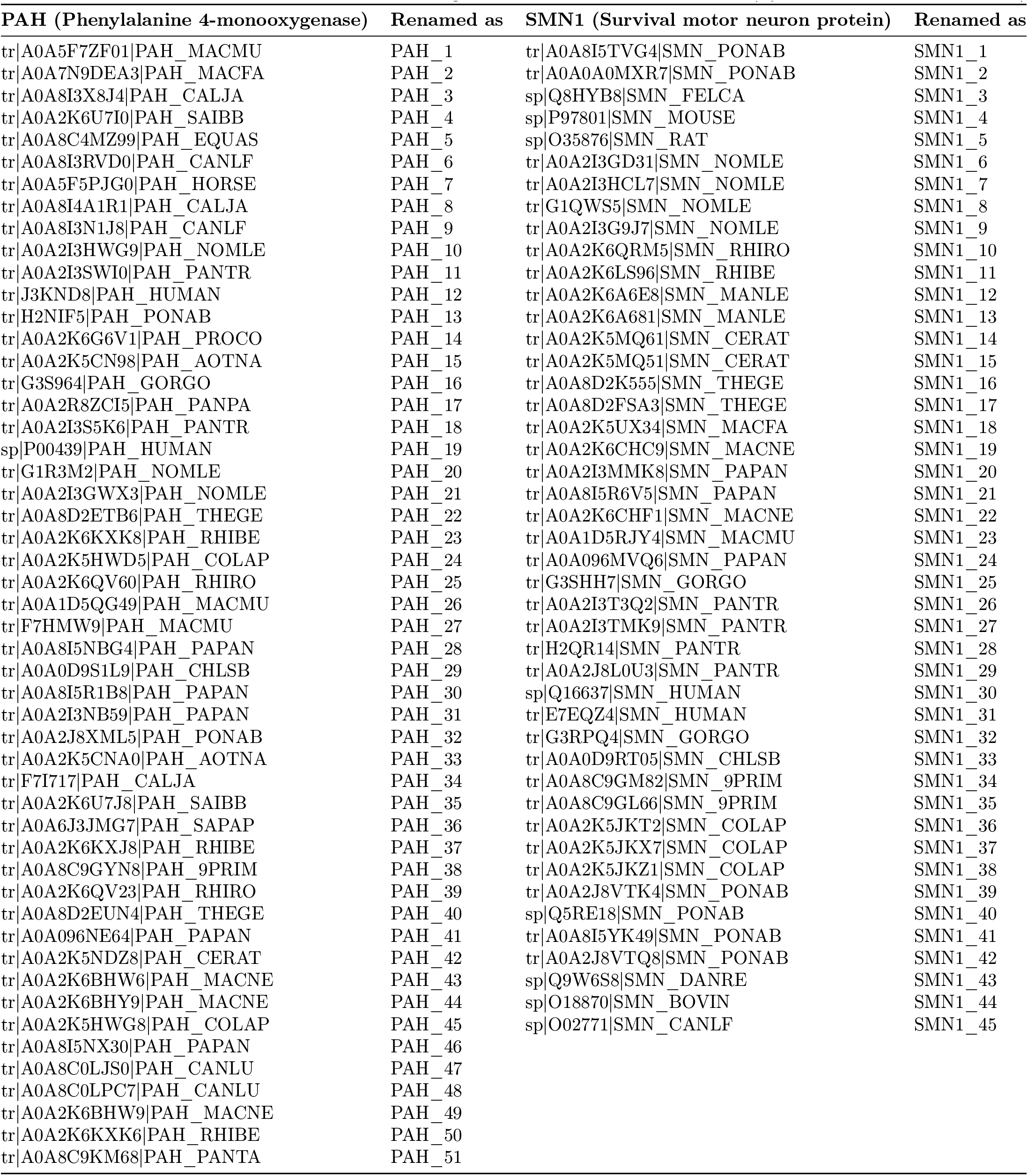
List of 51 PAH and 45 SMN1 proteins from various organisms with their associated Uniprot ID (Hyperlinked with respective Uniprot)

## 3. Methods

### 3.1. Composition profiler of PAH and SMN1 proteins

The Composition Profiler was used to generate an amino acid composition profile of all the PAH and SMN1 proteins analyzed in this study [49]. This set of amino acid sequences was the query set and the ‘Protein Data Bank Select 25’ was the background set. We also generated a composition profile for experimentally validated disordered proteins from the DisProt [50]. The generated profiles represent plots showing normalized enrichment or depletion of a given residue calculated as 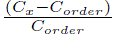, where *C_x_* is the content of a given residue in its query protein, and *C_order_* is the content of the same residue in the PDB Select 25.

### 3.2. Determining amino acid frequency composition of PAH and SMN1 proteins

The count of every amino acid in a sequence, termed amino acid frequency, was computed for all PAH and SMN1 proteins [51, 52, 53]. Furthermore, the percentage of amino acids in a sequence (obtained from dividing the amino acid frequencies by the length of that sequence and multiplied by 100) is termed the relative frequency of amino acids in that sequence. Relative frequency of 20 amino acids represents a 20-dimensional vector for each protein sequence.

#### 3.2.1. Evaluating Shannon entropy of PAH and SMN1 sequences

Shannon entropy (SE) is a measure of the information content in a system [54]. SE of each protein sequence is evaluated by the formula:

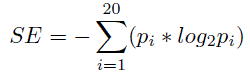

*p_i_* is the count of amino acid *i* in that sequence divided by the length of that sequence [51]. *SE* reflects the degree of randomness in the amino acid count in a given sequence. A higher value of *SE* indicates greater diversity, while a lower value indicates less diversity.

### 3.3. Amino acid frequency-based Shannon variability of position of PAH and SMN1 proteins

Shannon entropy is deployed to estimate the variability of amino acid residues at each residue position across all aligned PAH and SMN1 sequences, respectively. The Shannon variability (*H_v_*) for every position is defined as follows:

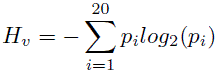

Here *p_i_*represents the fraction of residues of amino acid type *i* at a particular position. *H_v_*ranges from 0 (only one residue is present at that position) to 4.322 (all 20 residues are equally represented in that position) [55]. Typically, positions with *H_v_ >* 2 are considered *variable*, whereas those with *H_v_ <* 2 are considered *conserved*. *Highly conserved* positions are those with *H_v_ <* 1 [55].

By analyzing per-residue variability using Shannon entropy, one can identify functionally important residues within the receptor protein family. In addition, per-residue variability analysis using Shannon entropy can help identify potential drug targets within the receptor protein family. Residues that are highly variable among sequences may be more amenable to small molecule binding, as they may have more flexibility and plasticity in their binding pockets [56].

### 3.4. Determining the homogeneous poly-string frequency of amino acids in PAH and SMN1 proteins

A homogeneous poly-string of length *n* is defined as *n* consecutive occurrence of a particular amino acid [57, 58]. For example, *KKKWWKKWW* represents one homogeneous poly-string of K (Lys) with length 3, one homogeneous poly-string of K with length 2, and two homogeneous poly-strings of W (Trp) with length 2. Note that while counting the number of a homogeneous poly-string of length *n*, only the exclusive/exact occurrence of length *n* would be taken into consideration. The maximum lengths of homogeneous poly-strings considering all amino acids across all sequences were computed and accordingly, counts of homogeneous poly-strings of all possible lengths (starting from 1 to maximum length) for each amino acid present in a given protein sequence were enumerated [57].

### 3.5. Evaluating polar, non-polar residue profiles of PAH and SMN1 proteins

Every amino acid in a given protein sequence was identified as polar(P) or non-polar(Q). Thus, every protein sequence became a binary sequence with two symbols: P and Q. Through this binary P-Q profile, a spatial assembly of polar and non-polar residues over a protein sequence was depicted [57, 59, 60, 61].

#### 3.5.1. Change response sequences based on polar, non-polar residue profiles

There are four possible changes between two consecutive residues of a polar and non-polar profile, namely Polar to Polar (PP), Polar to Non-polar (PN), Non-polar to Non-Polar (NN), and Non-polar to Polar (NP) [57]. Such changes were accounted in the form of a sequence according to the spatial sequential arrangement of polar-non-polar residues in a given P-Q binary profile. We call this sequence “P-Q Change Response Sequence (*CRS_P Q_*)”. The frequency of each of the four changes was enumerated from the binary polar-non-polar profile corresponding to each PAH/SMN1 protein [57].

### 3.6. Evaluating acidic, basic, neutral residue profiles of PAH and SMN1 proteins

Every amino acid in a given PAH/SMN1 protein sequence was identified as acidic (A), basic (B), and neutral (N). Thus, every protein sequence became a ternary valued sequence (A-B-N profiles) with three symbols: A, B, and N [57].

#### 3.6.1. Change response sequences based on acidic-basic-neutral residue profiles

There are nine possible changes between two consecutive residues of an A-B-N profile, namely Acidic to Acidic (AA), Acidic to Basic (AB), Acidic to Neutral (AN), Basic to Acidic (BA), Basic to Basic (BB), Basic to Neutral (BN), Neutral to Acidic (NA), Neutral to Basic (NB), Neutral to Neutral (NN). Such changes were accounted in the form of a sequence according to the spatial sequential arrangement of acidic, basic, and neutral residues in a given A-B-N ternary profile. We designate this sequence “A-B-N Change Response Sequence (*CRS_ABN_*)”. The frequency of each of the nine changes was counted for a given ternary A-B-N profile corresponding to each PAH/SMN1 protein [57].

### 3.7. Evaluating intrinsic protein disorder of PAH and SMN1 proteins

Predisposition for intrinsic disorder of all the PAH and SMN1 proteins analyzed in this study was determined using a set of commonly used per-residue disorder predictors, such as PONDR® VLS2, PONDR® VL3, PONDR® VLXT, PONDR® FIT, IUPred-Long, IUPred-Short [62, 63, 64, 65, 66, 67]. A web platform called Rapid Intrinsic Disorder Analysis Online (RIDAO) was used to gather results from each predictor in bulk [68]. The percent of predicted intrinsically disorder residues (PPIDR) for each protein was used to classify each protein based on its level of disorder. A residue was considered to be disordered if it had a value of 0.5 or higher. Generally, a PPIDR value of less than 10% is taken to correspond to a highly ordered protein, PPIDR between 10% and 30% is ascribed to a moderately disordered protein, and PPIDR greater than 30% corresponds to a highly disordered protein [69, 70]. In addition to PPIDR, the mean disorder score (MDS) was calculated for each query protein as a protein length-normalized sum of all the per-residue disorder scores. The per-residue disorder score ranges from 0 to 1, where a score of 0 indicates fully ordered residues and a score of 1 indicates fully disordered residues. Residues with scores above the threshold of 0.5 were considered *disordered residues*. Residues with disorder scores between 0.25 and 0.5 were categorized as *highly flexible*, while those with scores between 0.1 and 0.25 were classified as *moderately flexible* [67]. We also utilized two binary predictors of disorder, the charge-hydropathy (CH) plot and the cumulative distribution function (CDF) (both accessible at Predictor of Natural Disordered Regions), to assess intrinsic disorder at the whole protein level [71, 72].

#### 3.7.1. Change response sequences based on intrinsic protein disorder residues

There are sixteen possible changes between two consecutive residues where residues were denoted as disordered (D), highly flexible (HF), moderately flexible (MF), and other (O) in PAH and SMN1 protein sequences namely disordered to disordered (D_D), disordered to highly flexible (D_HF), disordered to moderately flexible (D_MF), disordered to other (D_O) and similarly rest twelve HF_D, HF_HF HF_MF, HF_O, MF_D, MF_HF, MF_MF, MF_O, O_D, O_HF, O_MF, and O_O. Such changes were accounted for the form of a sequence according to the spatial sequential arrangement of D, HF, MF, and O residues for each protein sequence [57]. The frequency of each of the sixteen changes was counted from these change response sequences [57].

### 3.8. Structural and physicochemical features of PAH and SMN1 proteins

Structural and physicochemical descriptors extracted from sequence data have been widely used to characterize sequences and predict structural, functional, expression, and interaction profiles of proteins. iFeature (I-features) and ProtrWeb, two versatile Python-based toolkits were deployed to extract structural and physicochemical properties of the PAH and SMN1 proteins [73].

A list of 1162 features extracted using I-features, which were deployed in the present study was summarized in the following Table 3 [73].

**Table 3:**
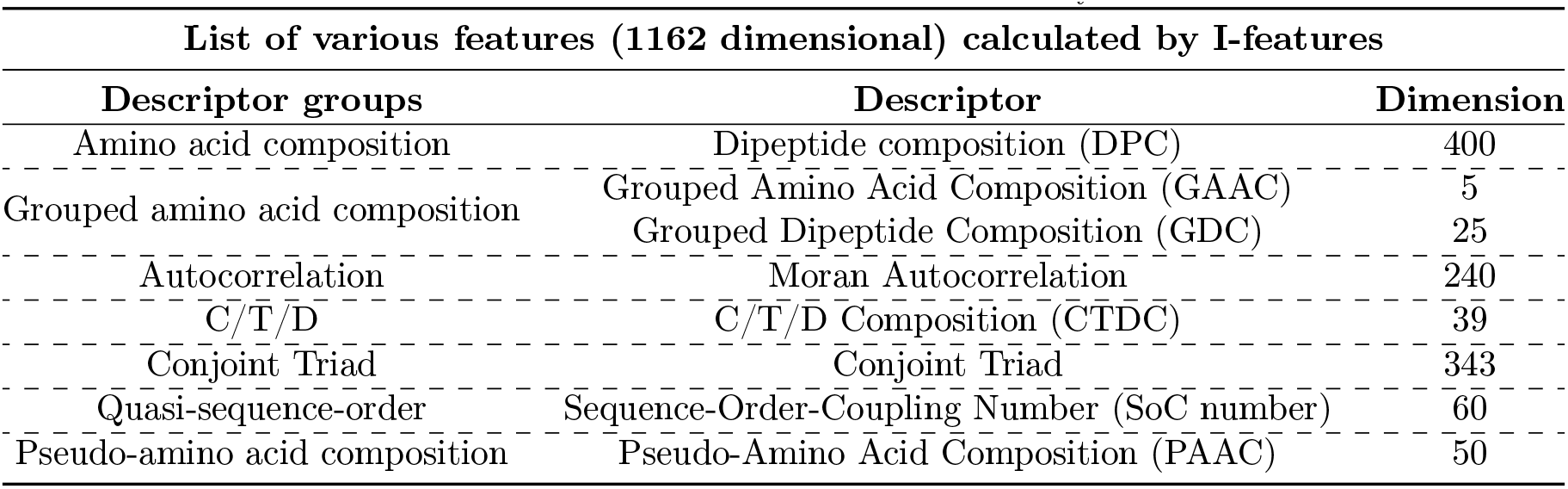
List of features of dimension 1162 calculated by I-features.

Another 8710 features extracted using ProtrWeb, which were deployed in the present study is summarized in Table 4 [73].

**Table 4:** List of various features (8710 dimensional) calculated by ProtrWeb.

| List of various features (8710 dimensional) calculated by ProtrWeb | Dimension |
| --- | --- |
| Tripeptide Composition | 8000 |
| Normalized Moreau-Broto Autocorrelation | 240 |
| Geary Autocorrelation | 240 |
| Quasi-Sequence-Order Descriptors | 100 |
| Pseudo-Amino Acid Composition | 50 |
| Amphiphilic Pseudo-Amino Acid Composition | 80 |

### 3.9. Formation of distance matrices and dendrograms

Euclidean distance was evaluated between feature vectors of all pairs of PAH/SMN1 protein sequences for each of the following six features: relative frequency of amino acids (dimension 20), the relative frequency of changes obtained from polar-nonpolar profiles (dimension 4), the relative frequency of changes obtained from acidic-basic-neutral profiles (dimension 9), the relative frequency of changes obtained from disordered, highly flexible, moderately flexible, and other residues (dimension 16), structural and physicochemical features (I-feature (dimension 1162) and ProtrWeb (dimension 8710)) [74]. Each of the structural and physicochemical features was normalized in the range of 0 to 100.

Sequence homology-based similarity matrices were obtained from Clustal Omega. Distance matrices were obtained by subtracting each entry of the similarity matrices from 100.

Each feature produces a distance matrix of dimension 51 *×* 51 (for PAH sequences)/45 *×* 45 (for SMN1 sequences). Different color thresholds (empirically chosen) were used in different dendrograms. If the color threshold had the value T, then each group of nodes whose linkage was less than T was assigned a unique color in the dendrogram and each color corresponds to a single cluster.

### 3.10. Agglomerated proximal sets of PAH and SMN1 proteins

A set of sequences ‘S’ (PAH/SMN1) is called an *‘agglomerated proximal set’* if every sequence of the set ‘S’ belongs to the same cluster for each of the seven dendrograms.

Mathematically, let {*c_i,_*_1_, *c_i,_*_2_, *c_i,_*_3_, *… c_i,k_*} be a set of *k* clusters corresponding to an aforementioned feature *f_i_*, where *i* = 1, 2, *…* 7. Note that *f*_1_ stand for the relative frequency of amino acids, *f*_2_ stands for sequence homology, *f*_3_ stands for the relative frequency of changes based on polar, non-polar residues, *f*_4_ stands for the relative frequency of changes based on acidic, basic, and neutral residues, *f*_5_ stands for change response based on intrinsic protein disordered regions, *f*_6_ stands for structural and physicochemical features derived from I-features, and *f*_7_ stands for structural and physico-chemical features derived from ProtrWeb.

A set *S* = {*s*_1_, *s*_2_, *s*_3_, *…, s_m_*_(_*_m≥_*_2)_} is said to be an ‘*agglomerated proximal set*’ if, for some *p ∈* {1, 2, *… k*}, *S ⊆ c_i,p_* for all *i* = 1, 2, *…* 7. Here, *s_j_* stands for a sequence of PAH/SMN1.

Following this definition, agglomerated proximal sets of PAH and SMN1 protein sequences were derived.

## 4. Results and analyses

### 4.1. Similitude and dissimilitude of PAH and SMN1 proteins based on amino acid frequency

#### 4.1.1. Compositional profile of PAH/SMN1 proteins

Characterized by noticeable differences, it has been shown that disordered proteins/regions are significantly depleted in bulky hydrophobic amino acid residues (I, L, and V) and aromatic amino acids (W, Y, F, and H), which are often involved in the formation of the hydrophobic core of a folded globular protein. Disordered proteins/regions also exhibit a low content of C, N, and M residues. These amino acids, namely C, W, I, Y, F, L, H, V, N, and M, which are depleted in disordered proteins and regions, are defined as order-promoting amino acids. On the other hand, disordered proteins and regions are substantially enriched in disorder-promoting amino acids, such as R, T, D, G, A, K, Q, S, E, and P [75, 76, 66, 77, 49]. These biases in the amino acid composition can be visualized using a web-based tool, the Composition Profiler, for the semi-automatic discovery of enrichment or depletion of amino acids in query proteins [49].

Amino acid composition profile analysis of 51 PAH sequences revealed that out of the ten order-promoting residues, four (W, V, N, and M) were significantly depleted in the PAH proteins, whereas four disorder-promoting residues (R, Q, E, and P) were significantly enriched (Figure 1).

**Figure 1:**
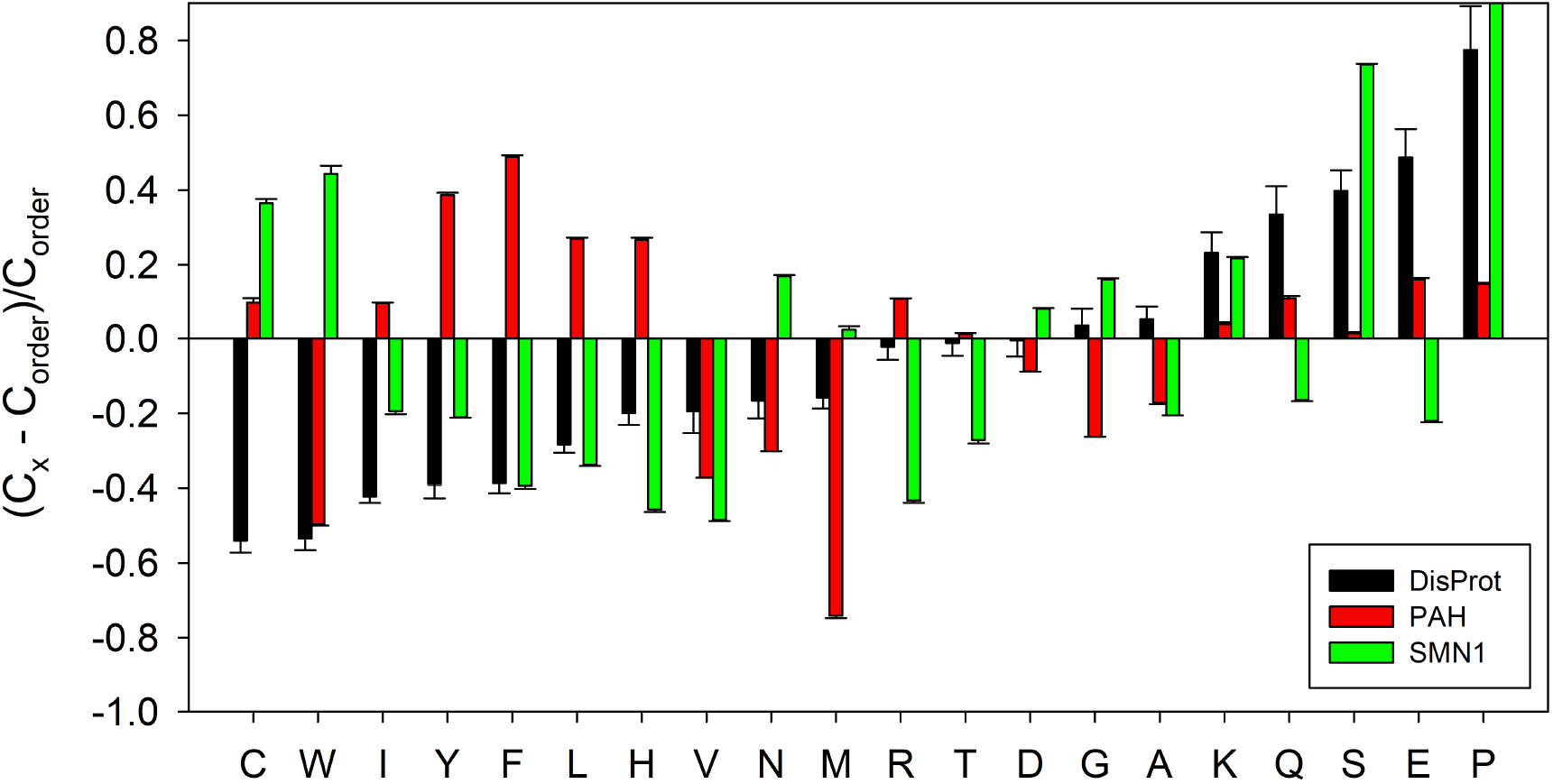
Amino acid composition profile of 51 PAH (red bars)and 45 SMN1 proteins (green bars). The fractional difference is calculated as 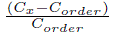, where *C_x_* is the content of a given amino acid in the query set (PAH/SMN1 proteins or known intrinsically disordered proteins), and *C_order_* is the content of a given amino acid in the background set (Protein Data Bank Select 25). The amino acid residues are ranked from most order-promoting residue to most disorder-promoting residue. Positive values indicate enrichment and negative values indicate depletion of a particular amino acid. The composition profile generated for experimentally validated disordered proteins from the DisProt database (black bars) is shown for comparison. In both cases, error bars correspond to standard deviations over 10,000 bootstrap iterations.

Analysis of the amino acid composition profile for 45 SMN1 sequences unveiled that, among the ten order-promoting residues, six (I, Y, F, L, H, and V) were significantly depleted in SMN1 proteins, while five disorder-promoting residues (D, G, K, S, and P) were significantly enriched (Figure 1).

Both PAH and SMN1 proteins exhibit a significant depletion of certain order-promoting residues, but the specific amino acids involved differ between the two proteins. Similarly, both proteins show a significant enrichment of disorder-promoting residues, again with differences in the specific amino acids affected.

#### 4.1.2. Relative frequency of amino acids and associated phylogenetic relationship

The analysis of relative amino acid frequencies in PAH and SMN1 sequences, depicted in histograms in Figure 2, provides valuable insights. Notably, Leu is consistently abundant in all PAH sequences, exceeding a remarkable 10% frequency. In contrast, Pro stands out as the most frequent amino acid in the majority of SMN1 sequences, except for two instances (SMN_25 and SMN_32, both from the GORGO group) where Ser takes the lead. Within PAH sequences, Met records the lowest frequency, except for four specific sequences (PAH_4, PAH_10, PAH_11, and PAH_12) where Trp assumes the lowest frequency position. Notably, the average frequencies of both Met and Trp in PAH sequences remain under 1 percent. In nearly all SMN1 sequences, His consistently displays the lowest frequency, with Met and Cys occupying the lowest frequency positions in SMN_1 and SMN_43, respectively.

**Figure 2:**
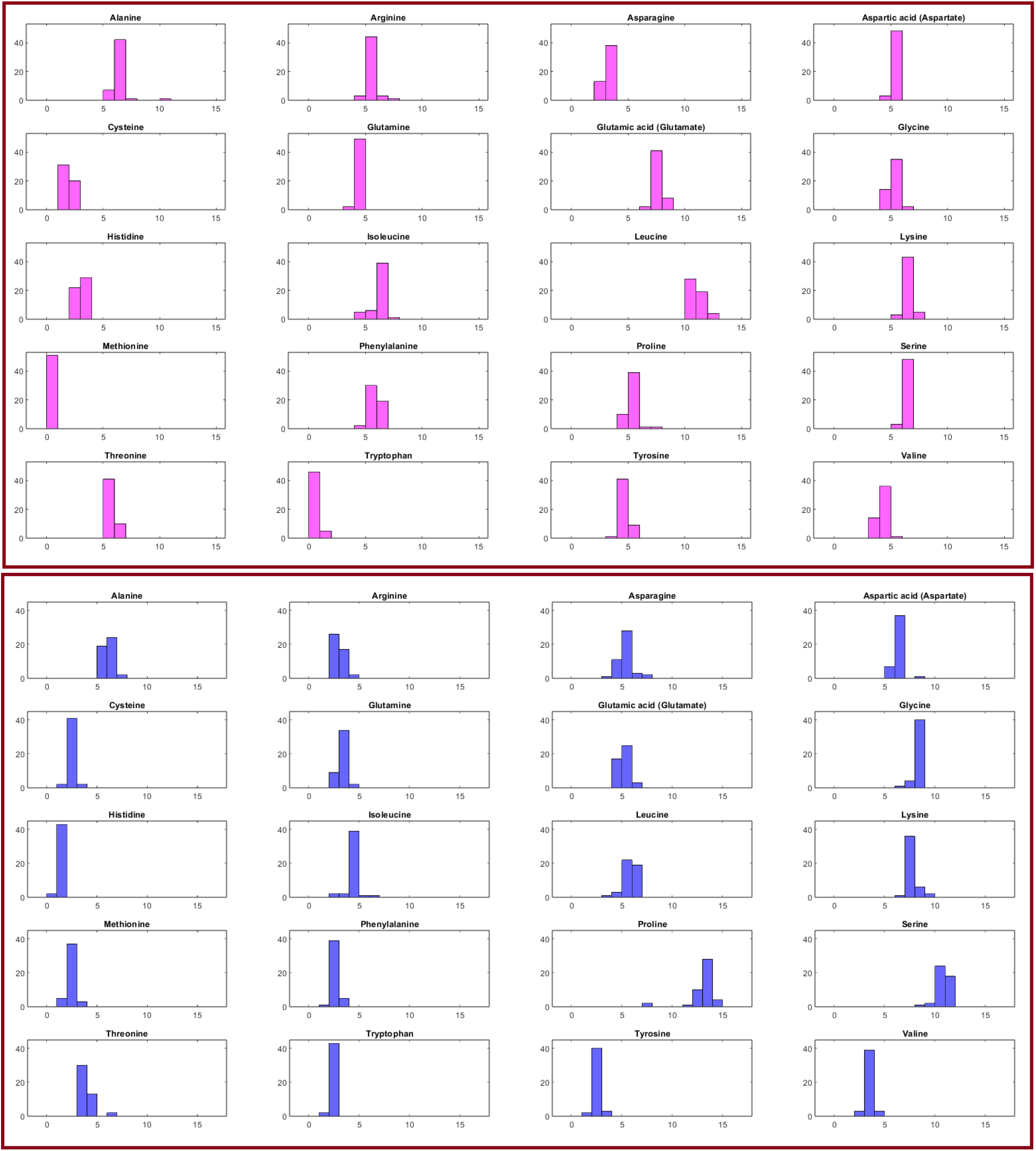
Histogram of the relative frequency of each amino acid in PAH proteins (Top), and SMN1 proteins (Bottom). X-axis denotes the percentage and Y-axis denotes the number of sequences.

These findings unveil distinctive amino acid composition patterns in PAH and SMN1 proteins, reflecting potential structural and functional implications. The prevalence of Leu in PAH sequences and the dominance of Pro in most SMN1 sequences underscore the significance of these amino acids in the respective proteins. Moreover, the variations in amino acid frequencies within specific sequences provide valuable insights into the diversity and unique characteristics of PAH and SMN1 proteins.

Utilizing a distance threshold of 1.3, a total of 51 PAH sequences were grouped into five distinct clusters, as illustrated in the dendrogram (Figure 3 (Top)). Furthermore, two PAH sequences, namely, PAH_3 and PAH_6 were distant from most the PAH sequences. Similarly, with a distance threshold of 1.87, 45 SMN1 sequences were partitioned into six clusters with SMN_43, SMN_25, and SMN_32 as outliers (Figure 3 (Bottom)). Notably, the largest clusters, referred to as Cluster-2 in both PAH and SMN1 proteins, consisted of 25 and 13 sequences, respectively, as highlighted in Table 5.

**Figure 3:**
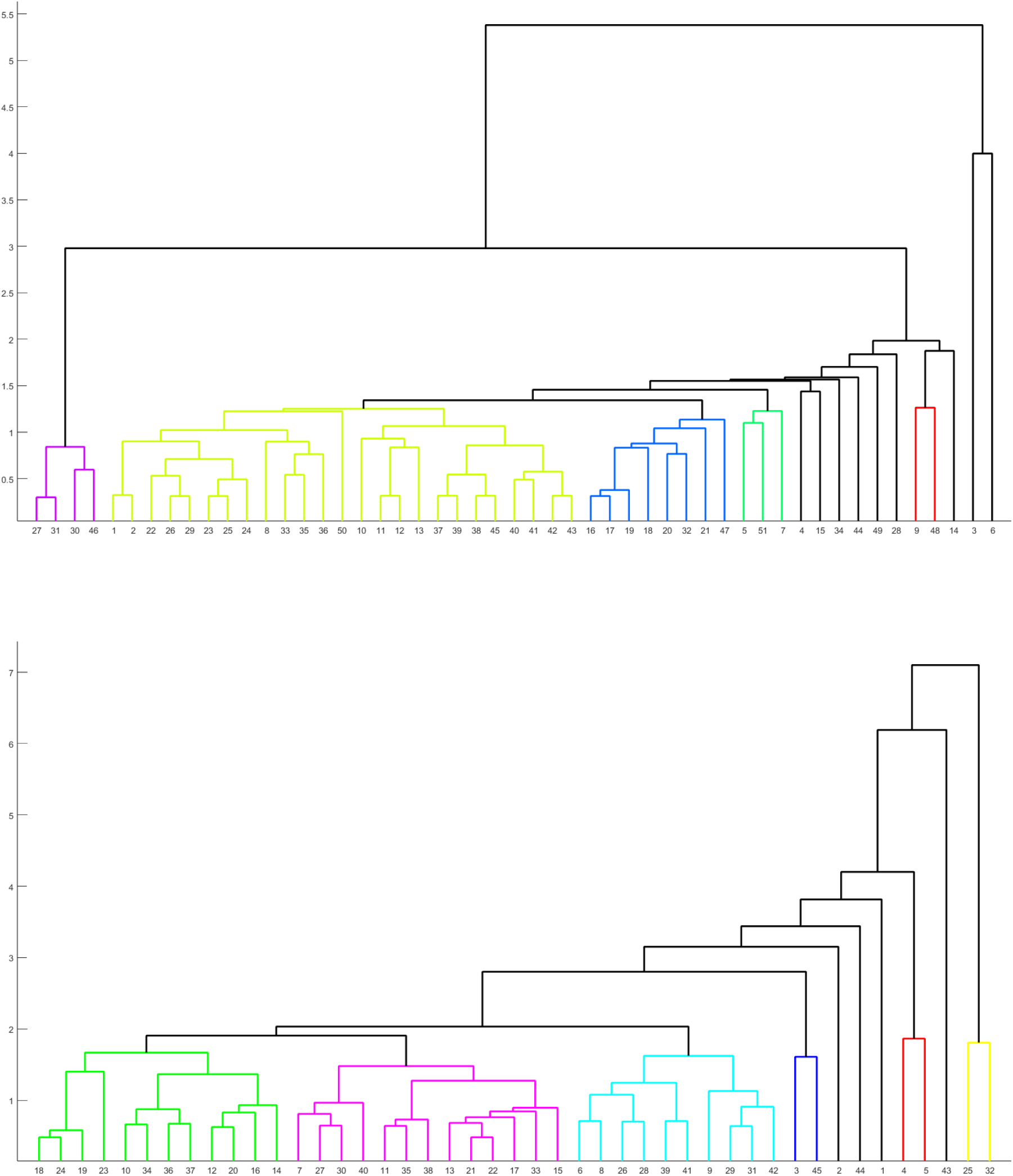
(Top): Phylogenetic relationship among the PAH proteins based on the relative frequency of amino acids. (Bottom): Phylogenetic relationship among the SMN1 proteins based on relative frequency of amino acids. Note that every color (other than black) in the phylogeny designates a cluster.

**Table 5:** Clusters derived from the relative frequency of amino acids in PAH and SMN1 proteins.

| Clusters | Sequences (PAH) |
| --- | --- |
| Cluster-1 | {27, 31, 30, 46} |
| Cluster-2 | {1, 2, 22, 26, 29, 23, 25, 24, 8, 33, 35, 36, 50, 10, 11, 12, 13, 37, 39, 38, 45, 40, 41, 42, 43} |
| Cluster-3 | {16, 17, 18, 19, 20, 32, 21, 47} |
| Cluster-4 | {5, 51, 7} |
| Cluster-5 | {9, 48} |
| Clusters | Sequences (SMN1) |
| Cluster-1 | {18, 24, 19, 23, 10, 34, 36, 37, 12, 20, 16, 14} |
| Cluster-2 | {7, 27, 30, 40, 11, 35, 38, 13, 21, 22, 17, 33, 15} |
| Cluster-3 | {6, 8, 26, 28, 39, 41, 9, 29, 31, 42} |
| Cluster-4 | {3, 45} |
| Cluster-5 | {4, 5} |
| Cluster-6 | {25, 32} |

#### 4.1.3. Shannon entropy of receptors

The Shannon entropy (SE) values for PAH and SMN1 sequences were grouped into six and five categories, respectively, based on their identical SEs, as detailed in Table 6. Furthermore, the SE values are notably clustered around 4.14 *±* 0.011 for PAH sequences and 4.08 *±* 0.011 for SMN1 sequences, as depicted in Figure 4. These observations indicate that the degree of disorderliness in amino acid frequencies is significantly high for both PAH and SMN1 sequences, as the SE values approach the maximum possible value of 4.322. This high level of disorderliness suggests that the amino acid composition within almost all PAH and SMN1 sequences is characterized by substantial variability and diversity, which may have implications for their structural and functional properties.

**Figure 4:**
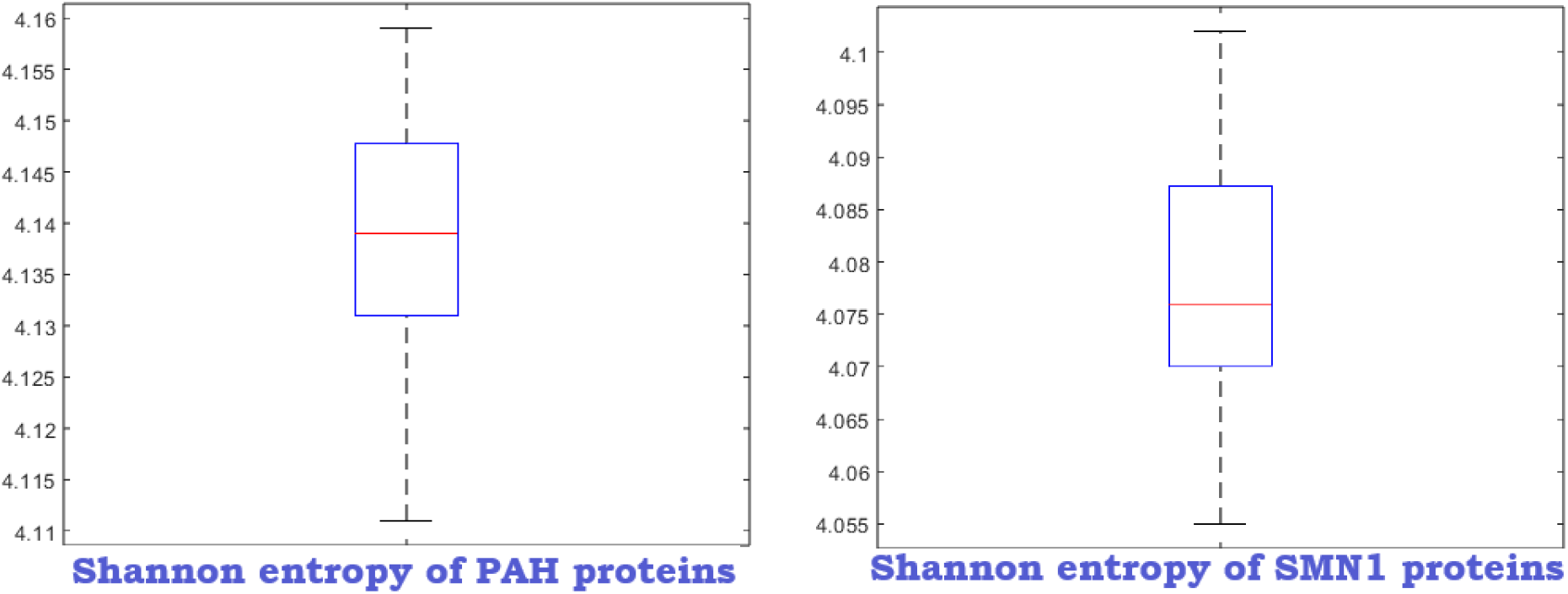
Box plot depicting the variation of Shannon entropy among the PAH and SMN1 protein sequences.

**Table 6:**
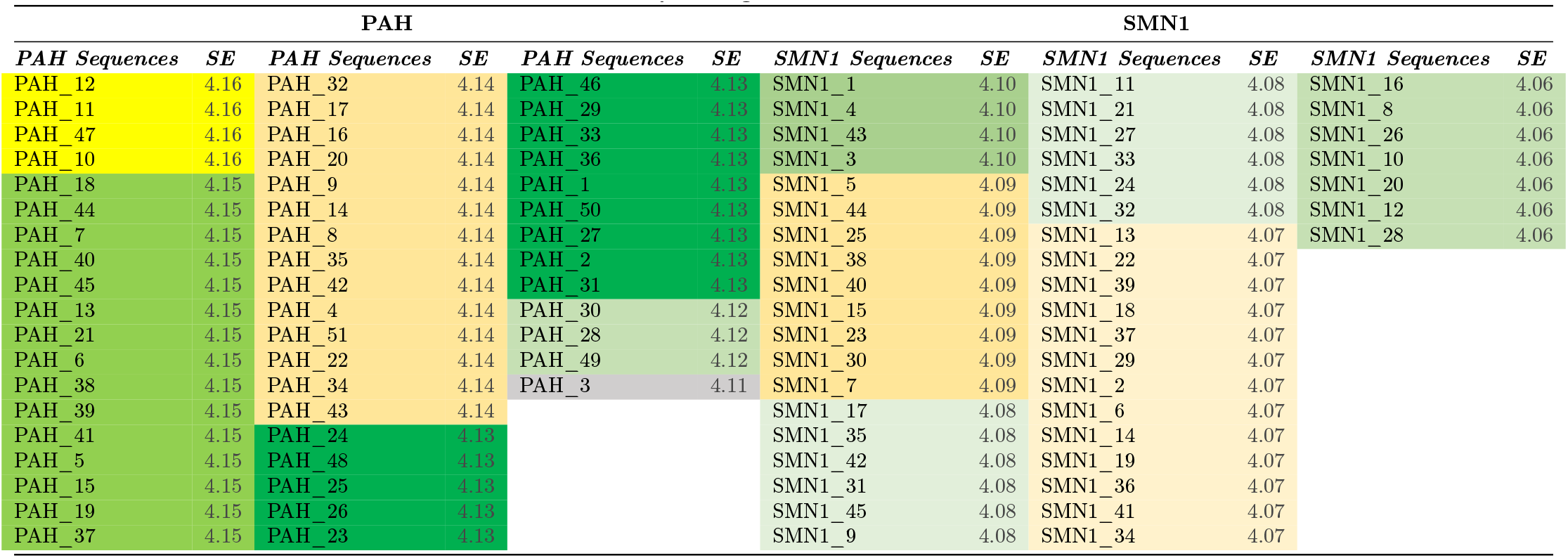
Shannon entropy among the PAH and SMN1 protein sequences.

### 4.2. Sequence homology and per-residue Shannon variability

#### 4.2.1. Sequence homology-based phylogenetic relationships and invariant residues

Using a distance threshold of 3.5, PAH sequences were grouped into five distinct clusters, depicted in Figure 5 (Up). PAH_6 stood out as an outlier. Simultaneously, SMN1 sequences were partitioned into three clusters and SMN_43 became an outlier, as shown in Figure 5 (Bottom). Notably, the most extensive clusters, identified as Cluster-2 for PAH and Cluster-3 for SMN1, contained 15 and 21 sequences, respectively, as detailed in Table 7.

**Figure 5:**
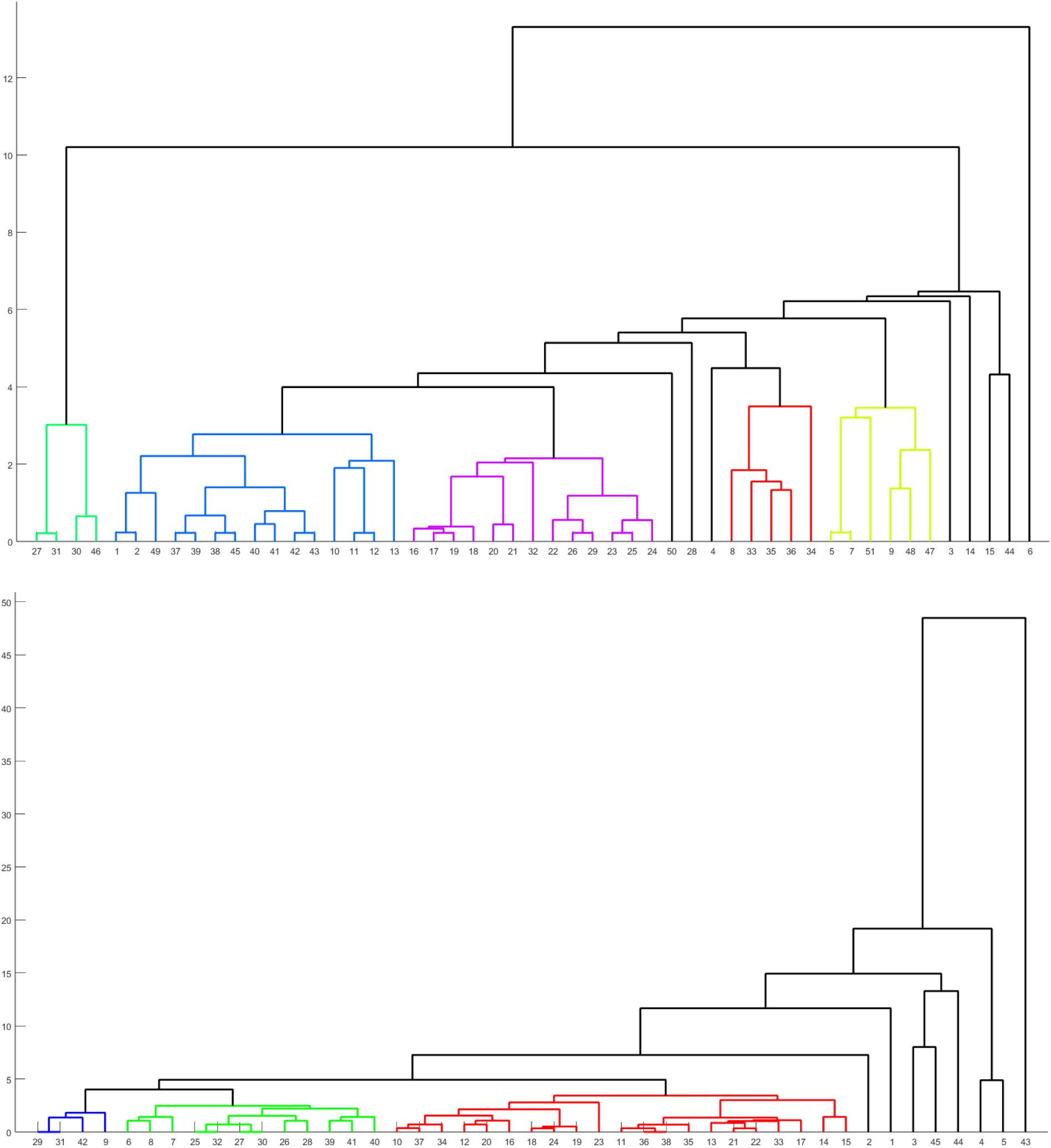
(Top): Phylogenetic relationship among the PAH proteins based on sequence homology. (Bottom): Phylogenetic relationship among the SMN1 proteins based on sequence homology. Note that every color (other than black) in the phylogeny designates a cluster.

**Table 7:** Clustering of PAH and SMN1 proteins based on sequence homology.

| Clusters | Sequences (PAH) |
| --- | --- |
| Cluster-1 | {27, 31, 30, 46} |
| Cluster-2 | {1, 2, 49, 37, 39, 38, 45, 40, 41, 42, 43, 10, 11, 12, 13} |
| Cluster-3 | {16, 17, 18, 19, 20, 21, 32, 22, 26, 29, 23, 25, 24} |
| Cluster-4 | {8, 33, 35, 36, 34} |
| Cluster-5 | {5, 7, 51, 9, 48, 47} |
| Clusters | Sequences (SMN1) |
| Cluster-1 | {29, 31, 42, 9} |
| Cluster-2 | {6, 8, 7, 25, 32, 27, 30, 26, 28, 39, 41, 40} |
| Cluster-3 | {10, 37, 34, 12, 20, 16, 18, 24, 19, 23, 11, 36, 38, 35, 13, 21, 22, 33, 17, 14, 15} |

Invariant residues in PAH(SMN1) sequences were determined through a multiple sequence alignment using Clustal Omega, referencing the tr|A0A8C9KM68|PAH_PANTA (sp|O02771|SMN_CANLF) sequence (see Table 8). We identified 23, 20, and 8 invariant residues of lengths 1, 2, and 3 in the PAH sequences, respectively. Notably, the longest invariant residue, spanning from amino acid position 88 to 136, had a length of 49 within the PAH sequences. Conversely, in the SMN1 sequences (Table 8), the longest invariant residue observed had a length of 5. It was noted that 66.2%, 63.8% and 2.4% residues were invariant in the regulatory N-terminal domain (amino acids 1-142), Catalytic central domain (amino acids 143-410), and oligomerization domain C-terminal amino acids 411-452), respectively.

**Table 8:** Invariant residues across PAH (SMN1) variants with reference to tr|A0A8C9KM68|PAH_PANTA (sp|O02771|SMN_CANLF)

| PAH |  |  |  |  |  | SMN1 |  |  |  |
| --- | --- | --- | --- | --- | --- | --- | --- | --- | --- |
| AA Position(s) | AA residues(s) | AA Position(s) | AA residues(s) | AA Position(s) | AA residues(s) | AA Position(s) | AA residues(s) | AA Position(s) | AA residues(s) |
| 20 | G | 148-150 | FAD | 327-342 | I-Q | 67-68 | RK | 134-137 | S-L |
| 30 | E | 152-153 | AY | 344-346 | CLS | 72-74 | KNK | 157 | E |
| 33 | G | 155 | Y | 349 | P | 87 | W | 161-162 | ST |
| 39 | L | 157-162 | H-P | 353-354 | PL | 90-91 | GD | 165 | S |
| 44-46 | END | 164-166 | VEY | 356-360 | L-A | 93 | C | 169 | S |
| 48-68 | N-T | 168-178 | E-F | 362 | Q | 97-101 | W-G | 186 | W |
| 70-71 | LD | 180-192 | T-E | 364 | Y | 104 | Y | 192-193 | PP |
| 73-74 | RS | 194-206 | N-F | 367-371 | T-P | 106-108 | ATI | 195 | P |
| 76 | P | 208 | E | 373-379 | Y-F | 110 | S |  |  |
| 78 | L | 210-215 | N-E | 381-387 | D-R | 112 | D |  |  |
| 80-85 | N-L | 217-222 | V-Q | 396-397 | PF | 117-120 | T-V |  |  |
| 88-136 | D-F | 224-260 | C-S | 404 | Y | 122-123 | YT |  |  |
| 138-142 | D-R | 262-290 | P-S | 406 | Q | 125-127 | YGN |  |  |
| 144-146 | RRK | 311-325 | I-D | 420 | L | 129-132 | E-N |  |  |

Invariant residues in PAH (SMN1) sequences across various organisms often correspond to critical functional regions of a protein [78]. These residues are involved in catalytic activity, ligand binding, and so on. As such, they can serve as signatures that define the sequence’s function. These invariant residues are conserved through evolution, because any mutation in these positions may disrupt the essential functions of the protein [78]. This conservation over time suggests the importance of these residues and can serve as a signature of evolutionary selection for that particular sequence. Invariant residues can be used as diagnostic markers in disease studies [79].

#### 4.2.2. Shannon variability of amino acid residue positions

The Shannon variability analyses revealed that an impressive 94.9% of residues in PAH variants and a substantial 90.88% of residues in SMN1 variants were highly conserved (as depicted in Figure 6). This high degree of conservation underscores the significance of these conserved residues in the structural and functional aspects of both PAH and SMN1 proteins. Such a level of conservation suggests that these residues likely play critical roles in maintaining the stability and functionality of these proteins, which could have implications for understanding their biological functions and the potential consequences of genetic variations in these conserved regions.

**Figure 6:**
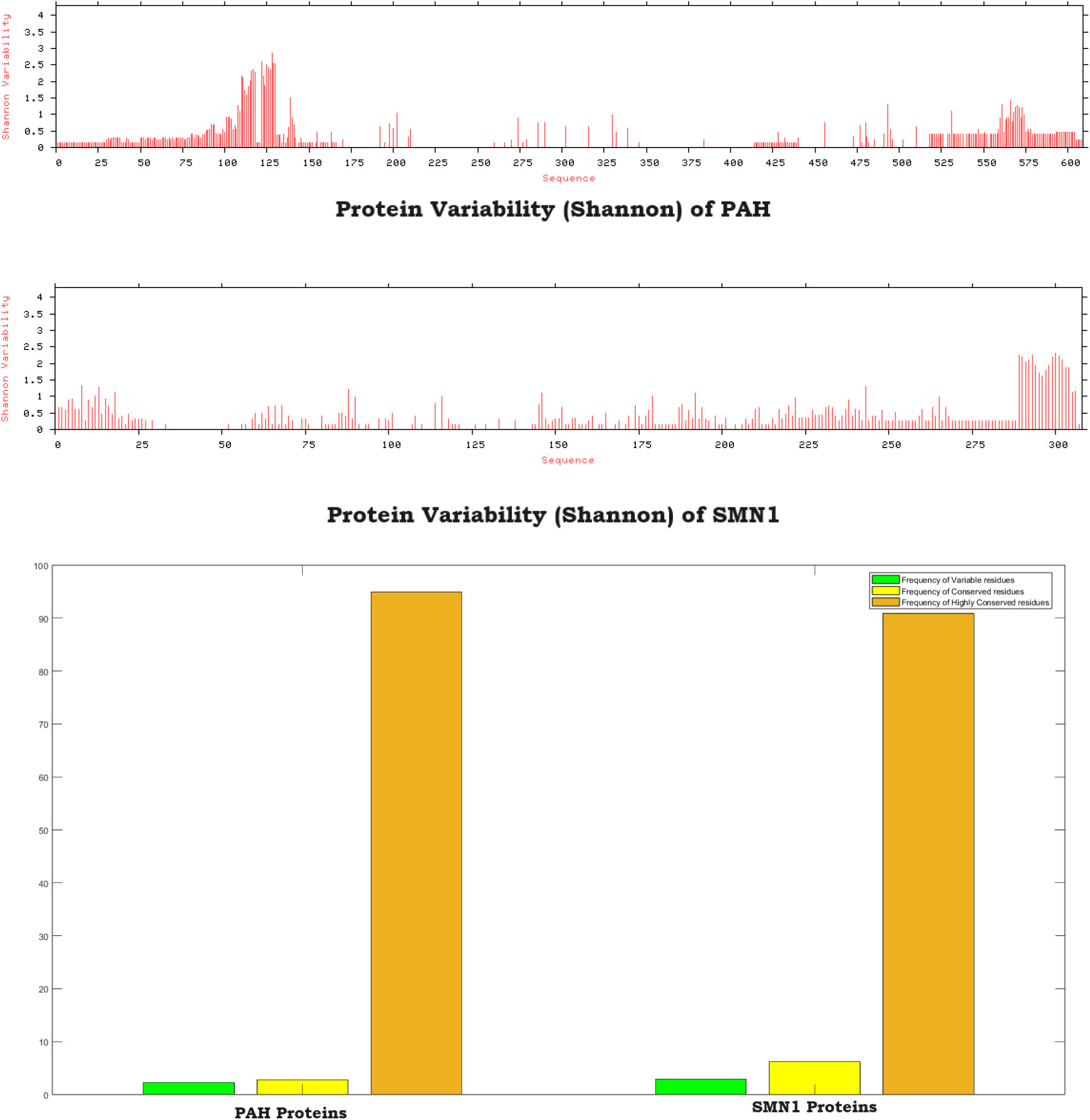
Shannon variability of amino acid residues in PAH and SMN1 proteins.

### 4.3. Homogeneous poly-string frequency of amino acids

The maximum length of a homogeneous poly-string was found to be 8 and 10 considering all amino acids across all PAH and SMN1 sequences, respectively. No poly-string of lengths 5, 6, or 7 was present in any of the 51 PAH sequences and no poly-string of length 7 was present in any of the 45 SMN1 sequences (Table 9). The frequencies of homogeneous poly-strings for each of the twenty amino acids separately are provided in **Supplementary file 1**.

**Table 9:**
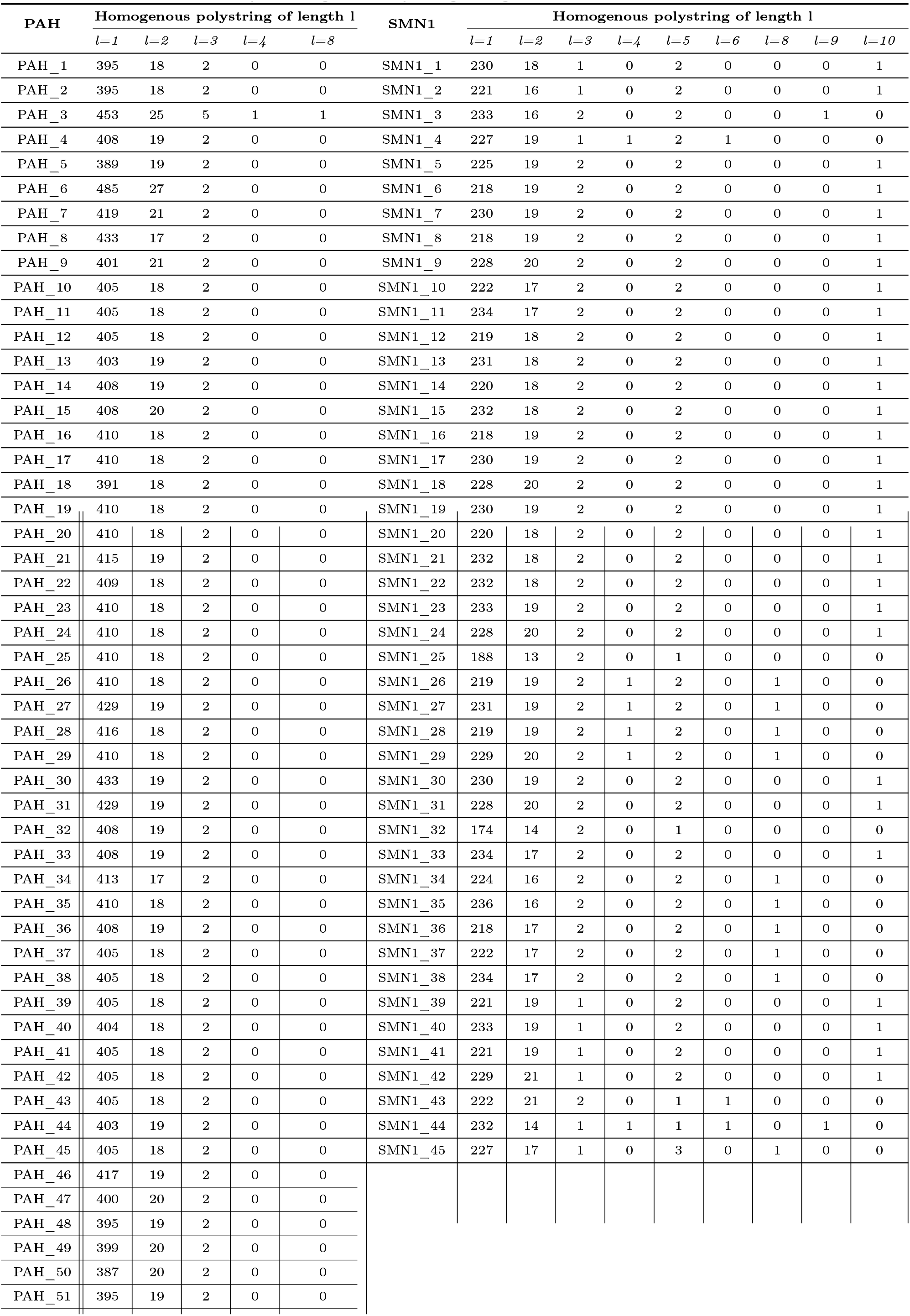
Frequency of homogeneous poly-string of length 1, 2, … 10 for each protein sequence.

| PAH | Homogenous polystring of length 1 |  |  |  |  | SMN1 | Homogenous polystring of length l |  |  |  |  |  |  |  |  |
| --- | --- | --- | --- | --- | --- | --- | --- | --- | --- | --- | --- | --- | --- | --- | --- |
|  | l=1 | l=2 | l=3 | l=4 | l=8 |  | l=1 | l=2 | l=3 | l=4 | l=5 | l=6 | l=8 | l=9 | l=10 |
| PAH_1 | 395 | 18 | 2 | 0 | 0 | SMN1_1 | 230 | 18 | 1 | 0 | 2 | 0 | 0 | 0 | 1 |
| PAH_2 | 395 | 18 | 2 | 0 | 0 | SMN1_2 | 221 | 16 | 1 | 0 | 2 | 0 | 0 | 0 | 1 |
| PAH_3 | 453 | 25 | 5 | 1 | 1 | SMN1_3 | 233 | 16 | 2 | 0 | 2 | 0 | 0 | 1 | 0 |
| PAH_4 | 408 | 19 | 2 | 0 | 0 | SMN1_4 | 227 | 19 | 1 | 1 | 2 | 1 | 0 | 0 | 0 |
| PAH_5 | 389 | 19 | 2 | 0 | 0 | SMN1_5 | 225 | 19 | 2 | 0 | 2 | 0 | 0 | 0 | 1 |
| PAH_6 | 485 | 27 | 2 | 0 | 0 | SMN1_6 | 218 | 19 | 2 | 0 | 2 | 0 | 0 | 0 | 1 |
| PAH_7 | 419 | 21 | 2 | 0 | 0 | SMN1_7 | 230 | 19 | 2 | 0 | 2 | 0 | 0 | 0 | 1 |
| PAH_8 | 433 | 17 | 2 | 0 | 0 | SMN1_8 | 218 | 19 | 2 | 0 | 2 | 0 | 0 | 0 | 1 |
| PAH_9 | 401 | 21 | 2 | 0 | 0 | SMN1_9 | 228 | 20 | 2 | 0 | 2 | 0 | 0 | 0 | 1 |
| PAH_10 | 405 | 18 | 2 | 0 | 0 | SMN1_10 | 222 | 17 | 2 | 0 | 2 | 0 | 0 | 0 | 1 |
| PAH_11 | 405 | 18 | 2 | 0 | 0 | SMN1_11 | 234 | 17 | 2 | 0 | 2 | 0 | 0 | 0 | 1 |
| PAH_12 | 405 | 18 | 2 | 0 | 0 | SMN1_12 | 219 | 18 | 2 | 0 | 2 | 0 | 0 | 0 | 1 |
| PAH_13 | 403 | 19 | 2 | 0 | 0 | SMN1_13 | 231 | 18 | 2 | 0 | 2 | 0 | 0 | 0 | 1 |
| PAH_14 | 408 | 19 | 2 | 0 | 0 | SMN1_14 | 220 | 18 | 2 | 0 | 2 | 0 | 0 | 0 | 1 |
| PAH_15 | 408 | 20 | 2 | 0 | 0 | SMN1_15 | 232 | 18 | 2 | 0 | 2 | 0 | 0 | 0 | 1 |
| PAH_16 | 410 | 18 | 2 | 0 | 0 | SMN1_16 | 218 | 19 | 2 | 0 | 2 | 0 | 0 | 0 | 1 |
| PAH_17 | 410 | 18 | 2 | 0 | 0 | SMN1_17 | 230 | 19 | 2 | 0 | 2 | 0 | 0 | 0 | 1 |
| PAH_18 | 391 | 18 | 2 | 0 | 0 | SMN1_18 | 228 | 20 | 2 | 0 | 2 | 0 | 0 | 0 | 1 |
| PAH_19 | 410 | 18 | 2 | 0 | 0 | SMN1_19 | 230 | 19 | 2 | 0 | 2 | 0 | 0 | 0 | 1 |
| PAH_20 | 410 | 18 | 2 | 0 | 0 | SMN1_20 | 220 | 18 | 2 | 0 | 2 | 0 | 0 | 0 | 1 |
| PAH_21 | 415 | 19 | 2 | 0 | 0 | SMN1_21 | 232 | 18 | 2 | 0 | 2 | 0 | 0 | 0 | 1 |
| PAH_22 | 409 | 18 | 2 | 0 | 0 | SMN1_22 | 232 | 18 | 2 | 0 | 2 | 0 | 0 | 0 | 1 |
| PAH_23 | 410 | 18 | 2 | 0 | 0 | SMN1_23 | 233 | 19 | 2 | 0 | 2 | 0 | 0 | 0 | 1 |
| PAH_24 | 410 | 18 | 2 | 0 | 0 | SMN1_24 | 228 | 20 | 2 | 0 | 2 | 0 | 0 | 0 | 1 |
| PAH_25 | 410 | 18 | 2 | 0 | 0 | SMN1_25 | 188 | 13 | 2 | 0 | 1 | 0 | 0 | 0 | 0 |
| PAH_26 | 410 | 18 | 2 | 0 | 0 | SMN1_26 | 219 | 19 | 2 | 1 | 2 | 0 | 1 | 0 | 0 |
| PAH_27 | 429 | 19 | 2 | 0 | 0 | SMN1_27 | 231 | 19 | 2 | 1 | 2 | 0 | 1 | 0 | 0 |
| PAH_28 | 416 | 18 | 2 | 0 | 0 | SMN1_28 | 219 | 19 | 2 | 1 | 2 | 0 | 1 | 0 | 0 |
| PAH_29 | 410 | 18 | 2 | 0 | 0 | SMN1_29 | 229 | 20 | 2 | 1 | 2 | 0 | 1 | 0 | 0 |
| PAH_30 | 433 | 19 | 2 | 0 | 0 | SMN1_30 | 230 | 19 | 2 | 0 | 2 | 0 | 0 | 0 | 1 |
| PAH_31 | 429 | 19 | 2 | 0 | 0 | SMN1_31 | 228 | 20 | 2 | 0 | 2 | 0 | 0 | 0 | 1 |
| PAH_32 | 408 | 19 | 2 | 0 | 0 | SMN1_32 | 174 | 14 | 2 | 0 | 1 | 0 | 0 | 0 | 0 |
| PAH_33 | 408 | 19 | 2 | 0 | 0 | SMN1_33 | 234 | 17 | 2 | 0 | 2 | 0 | 0 | 0 | 1 |
| PAH_34 | 413 | 17 | 2 | 0 | 0 | SMN1_34 | 224 | 16 | 2 | 0 | 2 | 0 | 1 | 0 | 0 |
| PAH_35 | 410 | 18 | 2 | 0 | 0 | SMN1_35 | 236 | 16 | 2 | 0 | 2 | 0 | 1 | 0 | 0 |
| PAH_36 | 408 | 19 | 2 | 0 | 0 | SMN1_36 | 218 | 17 | 2 | 0 | 2 | 0 | 1 | 0 | 0 |
| PAH_37 | 405 | 18 | 2 | 0 | 0 | SMN1_37 | 222 | 17 | 2 | 0 | 2 | 0 | 1 | 0 | 0 |
| PAH_38 | 405 | 18 | 2 | 0 | 0 | SMN1_38 | 234 | 17 | 2 | 0 | 2 | 0 | 1 | 0 | 0 |
| PAH_39 | 405 | 18 | 2 | 0 | 0 | SMN1_39 | 221 | 19 | 1 | 0 | 2 | 0 | 0 | 0 | 1 |
| PAH_40 | 404 | 18 | 2 | 0 | 0 | SMN1_40 | 233 | 19 | 1 | 0 | 2 | 0 | 0 | 0 | 1 |
| PAH_41 | 405 | 18 | 2 | 0 | 0 | SMN1_41 | 221 | 19 | 1 | 0 | 2 | 0 | 0 | 0 | 1 |
| PAH_42 | 405 | 18 | 2 | 0 | 0 | SMN1_42 | 229 | 21 | 1 | 0 | 2 | 0 | 0 | 0 | 1 |
| PAH_43 | 405 | 18 | 2 | 0 | 0 | SMN1_43 | 222 | 21 | 2 | 0 | 1 | 1 | 0 | 0 | 0 |
| PAH_44 | 403 | 19 | 2 | 0 | 0 | SMN1_44 | 232 | 14 | 1 | 1 | 1 | 1 | 0 | 1 | 0 |
| PAH_45 | 405 | 18 | 2 | 0 | 0 | SMN1_45 | 227 | 17 | 1 | 0 | 3 | 0 | 1 | 0 | 0 |
| PAH_46 | 417 | 19 | 2 | 0 | 0 |  |  |  |  |  |  |  |  |  |  |
| PAH_47 | 400 | 20 | 2 | 0 | 0 |  |  |  |  |  |  |  |  |  |  |
| PAH_48 | 395 | 19 | 2 | 0 | 0 |  |  |  |  |  |  |  |  |  |  |
| PAH_49 | 399 | 20 | 2 | 0 | 0 |  |  |  |  |  |  |  |  |  |  |
| PAH_50 | 387 | 20 | 2 | 0 | 0 |  |  |  |  |  |  |  |  |  |  |
| PAH_51 | 395 | 19 | 2 | 0 | 0 |  |  |  |  |  |  |  |  |  |  |

In the analysis of PAH sequences, a distinct pattern emerges regarding the lengths and amino acid compositions of homogeneous poly-strings. Among all PAH sequences, only PAH_3 had one poly-string of length 4 (of Arg) and another of length 8 (of Ala), whereas the remaining 50 PAH sequences exhibit poly-strings of maximum length 3 underscoring a consistent structural feature. Furthermore, it was noticed that only PAH_3 possessed five poly-strings (of Ala, Pro, Glx, and Lys) of length 3, while each of the remaining 50 PAH sequences contained two poly-strings (one of Glx and another of Lys) of length 3. Unique characteristic of PAH_3 suggests a potential functional significance of this particular sequence. None of the PAH had a poly-string of length 2 consisting of Cis, His or Val except PAH_47, PAH_21, and PAH_14, which had one poly-string of length 2 of Cys, His, and Val, respectively. None of the PAH had poly-string of length 2 containing Asp or Met or Asn or Trp. This may imply a selective constraint or functional constraint in the amino acid composition of these poly-strings. All 51 PAH sequences contained a single occurrence of FF, II, and YY (poly-strings of Phe, Ile, and Tyr with length 2). KK (poly-strings of Lys with length 2) was present with frequency 2 in all PAH, except PAH_49 which had frequency 3 of the same. These findings showcase a consistent pattern across most sequences.

Turning attention to the analysis of SMN1 sequences, a striking dominance of Pro in poly-strings of various lengths is evident. 29 out of 45 SMN1 sequences had single poly-string of length 10 and all of them comprised of proline. Poly-string of length 9 appeared in SMN1_3 and SMN1_44 only (with frequency 1 consisting of proline) and in no other SMN1. Additionally, 10 out of 45 SMN1 sequences had single poly-string of length 8, and all of them were composed of Pro. Sequences that had poly-string of length 10 did not have poly-strings of length 8. SMN1_4, SMN1_43, and SMN1_44 only had poly-string of length 6 (with frequency 1 consisting of Pro, Pro, and Gly, respectively) and no other SMN1 displayed poly-string of length 6. Except for SMN1_45, all SMN1 possessed poly-strings of length 5 with frequency 1 or 2 and made of Pro. Along with Pro, SMN1_45 had another poly-string of length 5 consisting of Gly. Out of 45, only 6 SMN1 sequences (SMN1_4, SMN1_26, SMN1_27, SMN1_28, SMN1_29, and SMN1_44) had poly-string of length 4 (all with frequency 1 and composed of Pro). Single occurrences of VVV (poly-string of Val with length 3) were noticed in all sequences except SMN1_43 in which VVV was absent. SMN1_43 had two poly-strings (one of Ala and the other of Glx) of length 3 and among the rest 44 SMN1 sequences, 35 had a single occurrence of GGG (poly-string of Gly with length 3) along with VVV. WW and VV (poly-string of Trp and Val with length 2) were not identified in any of the 45 SMN1 sequences except SMN1_43 which possessed VV with frequency 2. The unique features in SMN1_43 imply sequence-specific variations that may contribute to distinct structural or functional properties. All SMN1 sequences had a single occurrence of EE and AA (poly-string of Glx and Ala with length 2) underscoring shared structural elements among all SMN1 sequences.

### 4.4. Polar, non-polar residue profiles of PAH/SMN1 proteins

The percentage distributions of polar and non-polar residues were computed for each PAH Figure 7(Top) and SMN1 Figure 7 (Bottom) sequence. The analysis revealed that the average ratio of polar to non-polar residues in PAH sequences was 1.11 *±* 0.03 and ranged from 0.99 to 1.18. The average ratio of polar to non-polar residues in SMN1 sequences was 0.99 *±* 0.072 and ranged from 0.91 to 1.29. It was noteworthy that the number of polar and non-polar residues was almost identical in each PAH and SMN1 sequence. This uniformity in the frequency of polar and non-polar residues underscores a specific compositional characteristic shared among these sequences.

**Figure 7:**
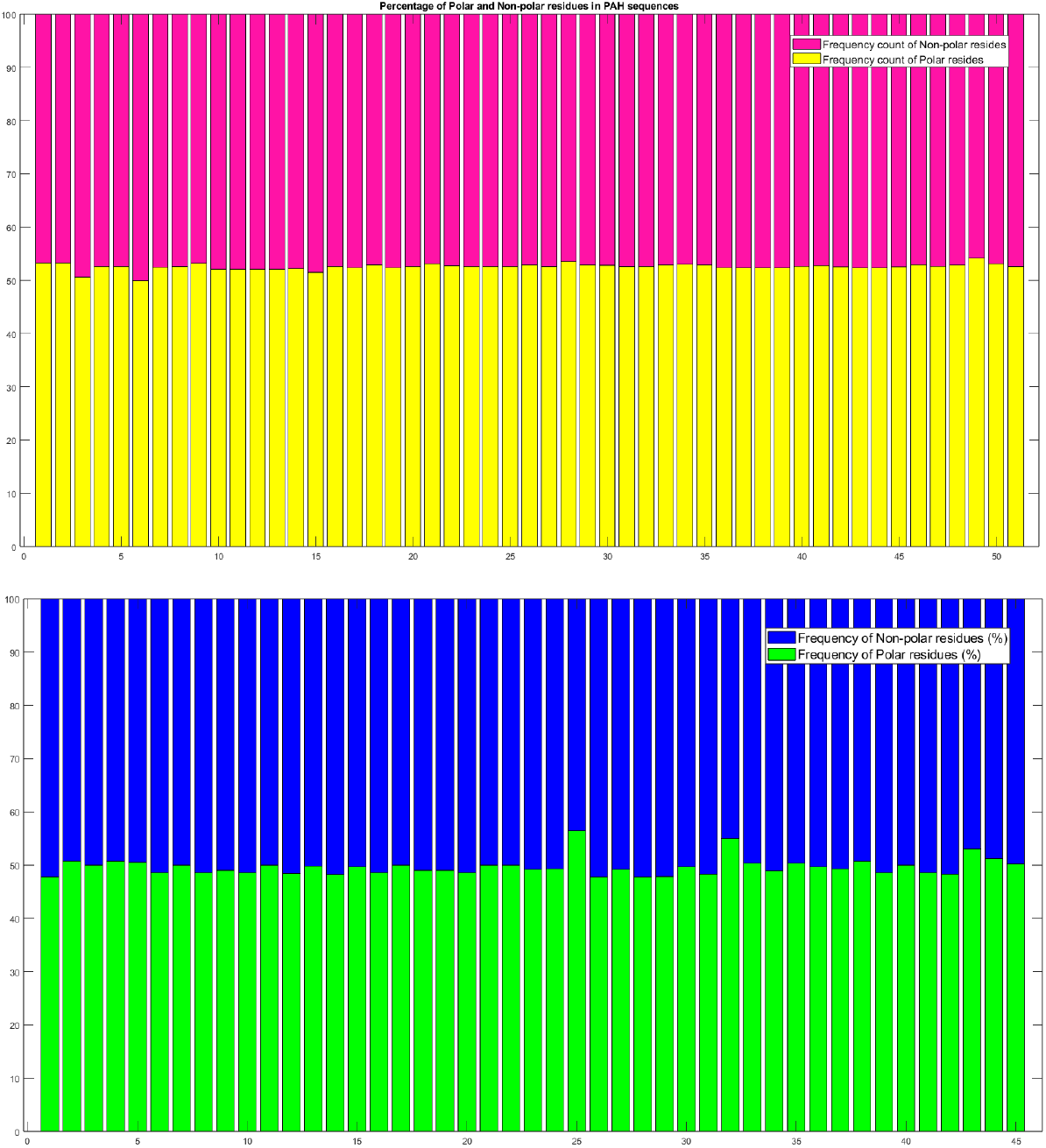
Percentages of polar, non-polar residues in PAH (Left) and SMN1 (Right) proteins.

#### 4.4.1. Change response sequences of polar, and non-polar profiles

Figure 8 (Left) illustrates the relative frequency distribution of four distinct changes: ‘PN’, ‘NP’, ‘PP’, and ‘NN’ within the PAH sequences. Notably, the median percentage of ‘PP’ was the highest among the four categories as PP for all PAH sequences range from 27.15% to 30.18%, except PAH_6. It’s interesting to observe that among these sequences, PAH_28 exhibited the lowest percentage of ‘NN’ changes at 21.88%, while two sequences, PAH_6 and PAH_3, displayed significantly high percentages of ‘NN’ changes (exceeding 25%) implying distinct functional or structural features in these sequences. The majority of PAH sequences exhibited a high percentage of polar-to-polar residue changes, indicating a propensity for the conservation of polar interactions within these sequences.

**Figure 8:**
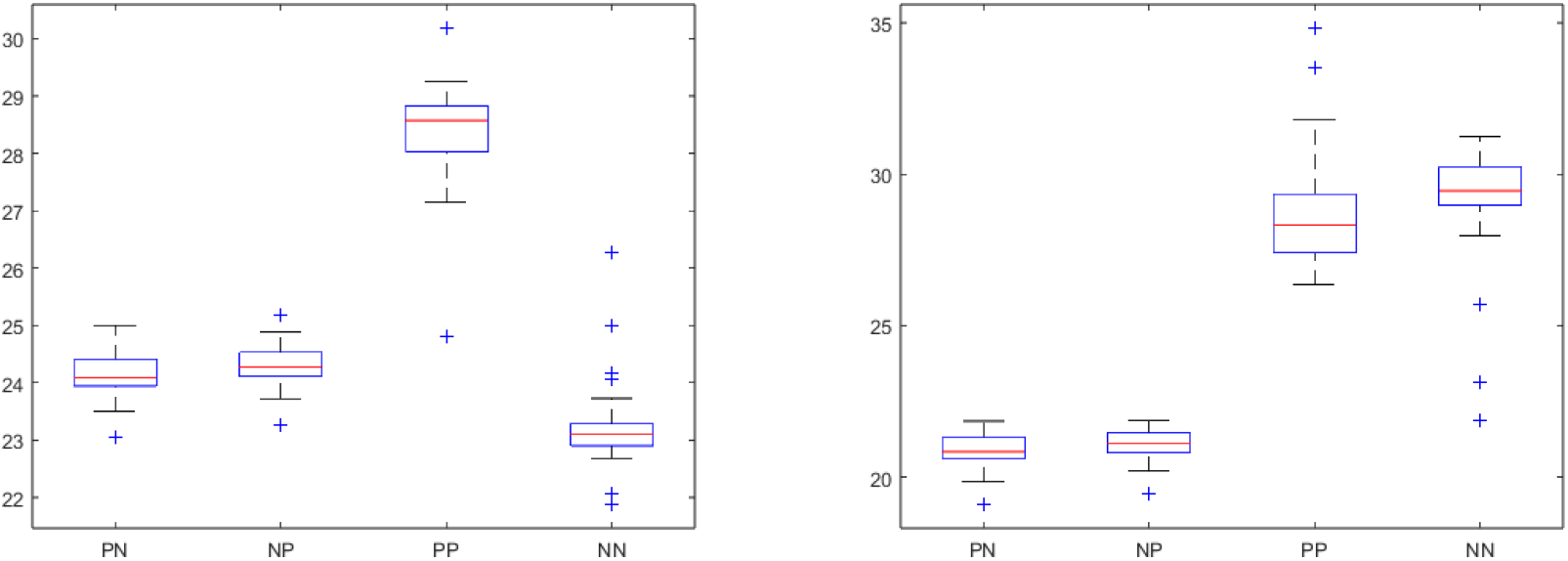
Box-plot of the relative frequency of PN, NP, PP, and NN changes in PAH (Top) and SMN1 (Bottom) proteins.

On the other hand, in the case of SMN1 sequences, the relative frequency distribution of the same four changes (‘PN’, ‘NP’, ‘PP’, and ‘NN’) reveals that the median percentage of ‘NN’ was the highest indicating a preference for conserving non-polar interactions within this set (see Figure 8 (Right)). SMN1_25 and SMN1_32 showed the lowest ‘NN’ changes at 21.88% and 23.11%, respectively, possibly reflecting a specific evolutionary adaptation in these sequences. Additionally, percentages of ‘PN’ and ‘NP’ changes were comparatively low for two sequences, SMN1_5 and SMN1_4, as indicated by their outlier status in the box plot (see Figure 8 (Right)).

The analyses of residue changes provide insights into the conservation and variation of specific interactions within the PAH and SMN1 sequences. These observations can be valuable for understanding the functional and structural characteristics of these sequences and may guide further investigations into their biological significance.

Based on a distance threshold of 0.7, 51 PAH sequences formed three clusters (Figure 9 (Up)), and 45 SMN1 sequences formed four clusters (Figure 9 (Bottom)) based on the distance threshold of 1.5. The largest clusters (cluster-2 for PAH and cluster-3 for SMN1) had 23 and 14 sequences, respectively, as noted in Table 10. PAH_3 and PAH_6 turned out to be distant from the rest of the PAH sequences while SMN_25, SMN_32, and SMN_43 became outliers among SMN1 sequences.

**Figure 9:**
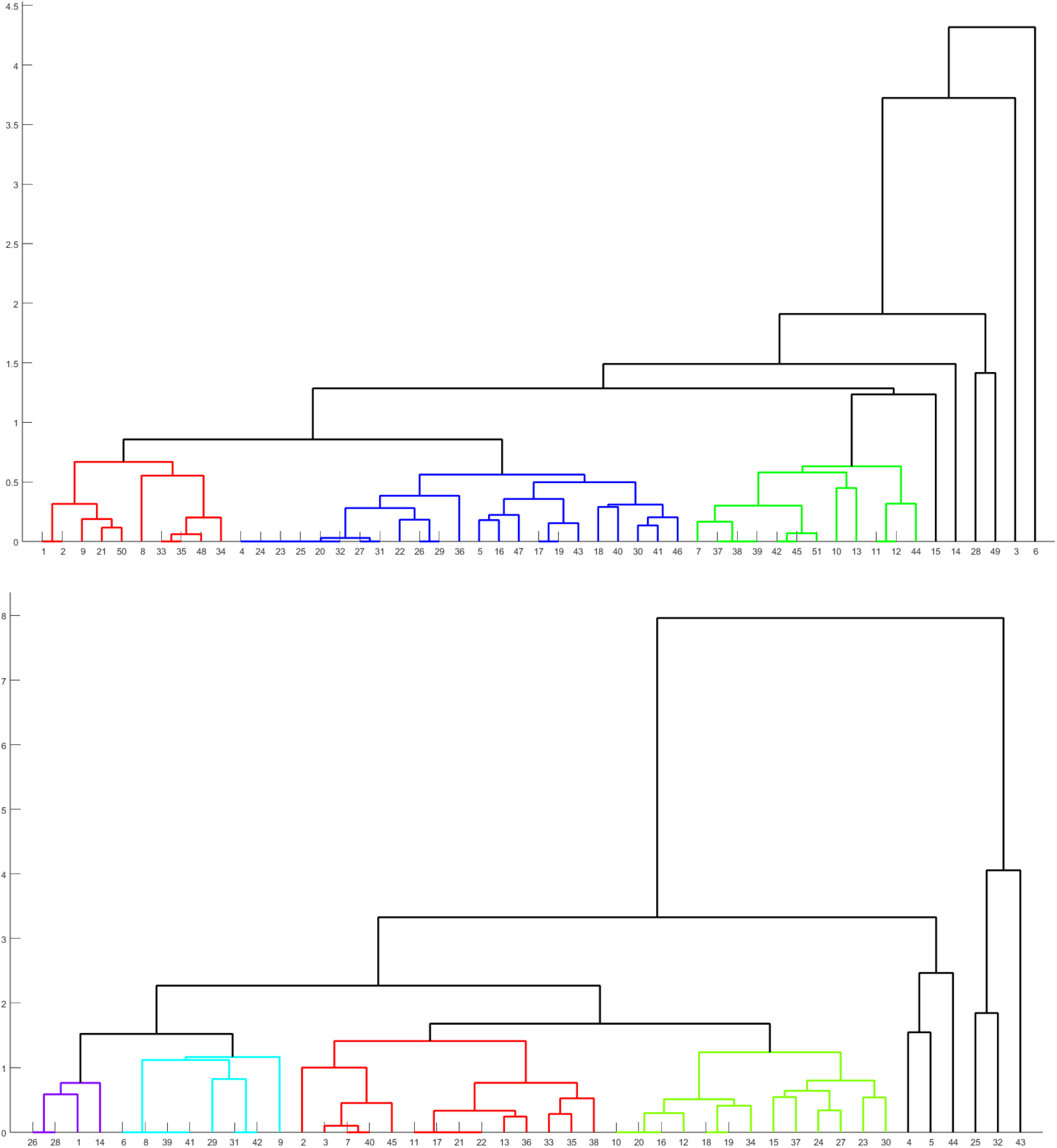
Phylogenetic relationship among the PAH (Top) and SMN1 proteins (Bottom) based on the relative frequency of PP, NP, PP, and NN changes as obtained from polar, non-polar profiles.

**Table 10:**
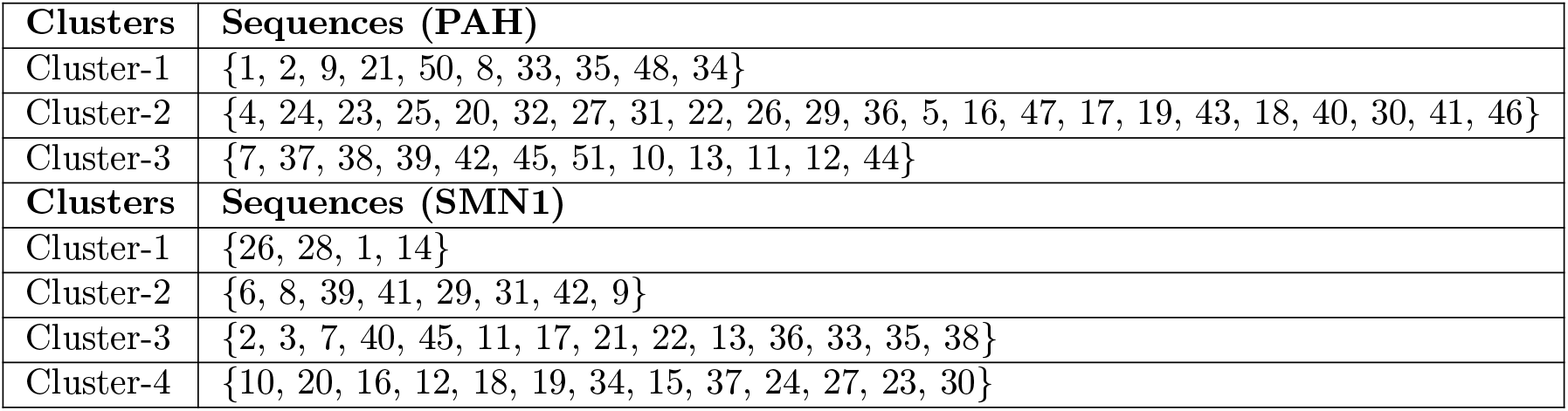
Clusters of PAH and SMN1 proteins based on the relative frequency of ‘PN’, ‘NP’, ‘PP’, and ‘NN’ changes as obtained from Polar, non-polar profiles.

### 4.5. Acidic, basic, neutral residue-based phylogenetic relationship

Based on the data presented in Figure 10, the analysis involved the calculation of the percentages of acidic, basic, and neutral residues for each PAH and SMN1 protein. This analysis encompassed 51 PAH sequences and 45 SMN1 sequences, revealing varying percentages of neutral residues ranging from 73.48% to 76.88% for PAH and from 72.59% to 79.59% for SMN1. The calculated ratio of acidic to basic residue percentages within the PAH sequences was determined to be 1.08 *±* 0.06, while for SMN1 sequences, it was 1.09 *±* 0.071. This indicates a relatively balanced distribution between acidic and basic residues in both PAH and SMN1 sequences.

**Figure 10:**
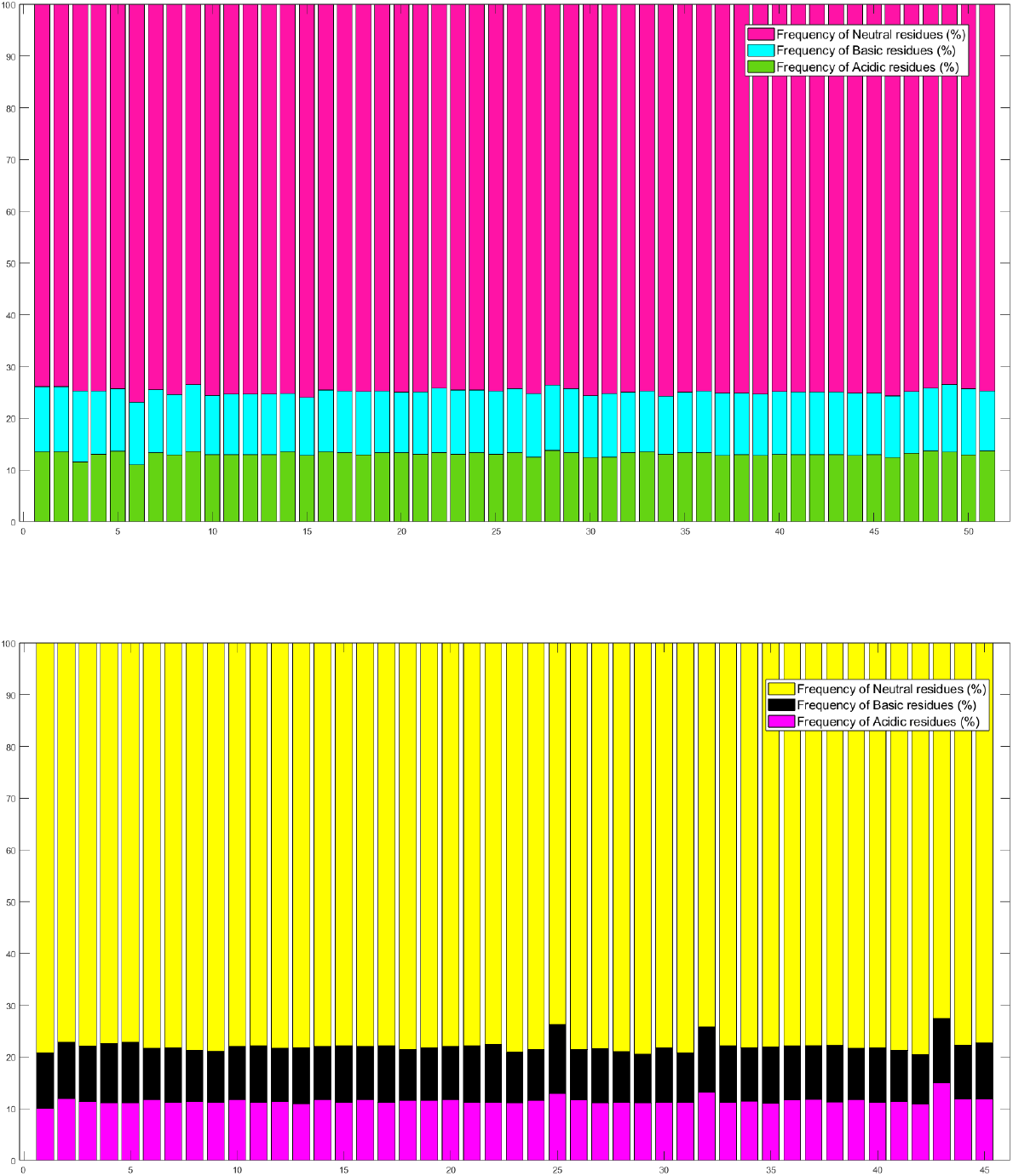
Percentages of acidic, basic, neutral residues in PAH (Top) and SMN1 (Bottom) proteins.

#### 4.5.1. Change response sequences of acidic, basic, and neutral profiles

The relative frequency distribution of nine different residue changes in PAH (SMN1) sequences, denoted as BA, NA, AA, BB, NB, AB, BN, NN, and AN, was visualized in Figure 11 (Left (Right)). It was found that the percentage of ‘NN’ change of residues in all PAH (SMN1) sequences was high in comparison to the other eight changes with values ranging from 55.36% to 59.56% (from 54.9% to 64.8%). Furthermore, it was observed that the percentage of acidic to basic residue changes was lowest in all SMN1 sequences, whereas the percentage of basic to basic (basic to acidic in some sequences) was found to be the minimum among all.

**Figure 11:**
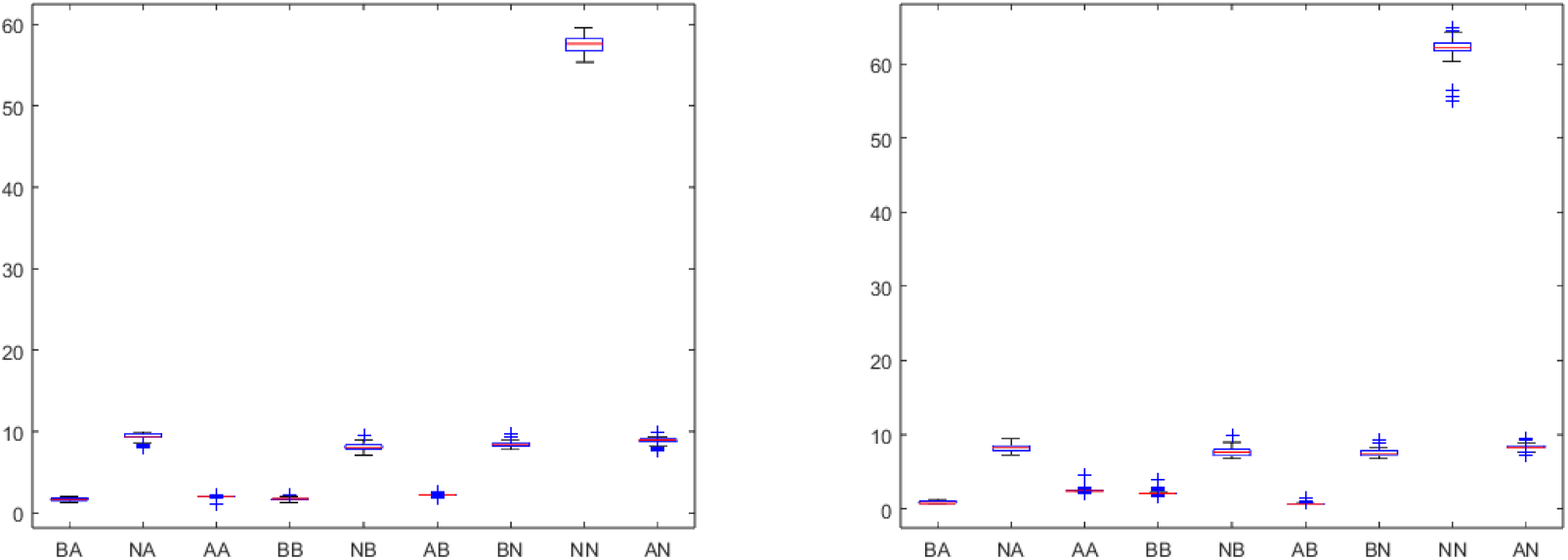
Box-plot of the relative frequency of all nine changes in PAH (Left) and SMN1 (Right) proteins.

Under a distance threshold of 1.25 (2), the analysis led to the formation of five distinct clusters from PAH and three from SMN1 sequences. This information is visually depicted in the dendrogram (Figure 12 (Top)) and summarized in Table 11. Notably, the largest cluster for PAH contained 21 sequences, as listed as Cluster-3 in Table 11. Among the three clusters, cluster-1 and cluster-3 contain the highest (33) and lowest (2) number of SMN1 sequences, respectively. Similar to previous dendrograms, PAH_3 and PAH_6 became outliers among PAH sequences, while SMN_25, SMN_32, and SMN_43 were found to be distant from the rest of the SMN1 sequences.

**Figure 12:**
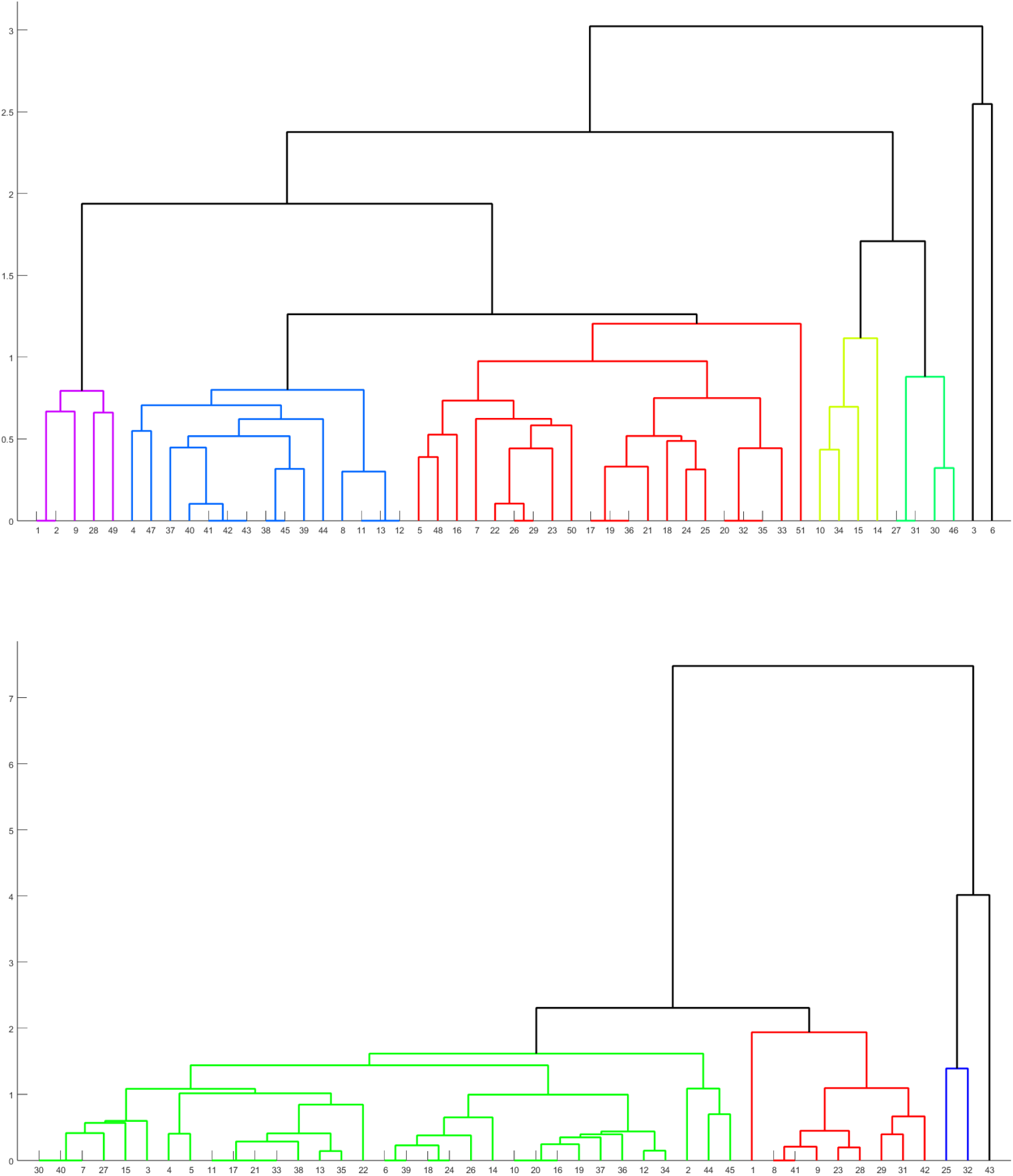
Phylogenetic relationship among the PAH (Top) and SMN1 (Bottom) proteins based on the relative frequency of BA, NA, AA, BB, NB, AB, BN, NN, and AN changes as obtained from acidic, basic, and neutral profiles.

**Table 11:** Clusters of PAH and SMN1 proteins based on the relative frequency of BA, NA, AA, BB, NB, AB, BN, NN, and AN changes as obtained from acidic, basic, and neutral profiles.

| Clusters | Sequences (PAH) |
| --- | --- |
| Cluster-1 | {1, 2, 9, 28, 49} |
| Cluster-2 | {4, 47, 37, 40, 42, 43, 38, 45, 39, 44, 8, 11, 13, 12} |
| Cluster-3 | {5, 48, 16, 7, 22, 26, 29, 23, 50, 17, 19, 36, 21, 18, 24, 25, 20, 32, 35, 33, 51} |
| Cluster-4 | {10, 34, 15, 14} |
| Cluster-5 | {27, 31, 30, 46} |
| Clusters | Sequences (SMN1) |
| Cluster-1 | {30, 40, 7, 27, 15, 3, 4, 5, 11, 17, 21, 33, 38, 13, 35, 22, 6, 39, 18, 24, 26, 14, 10, 20, 16, 19, 37, 36, 12, 34, 2, 44, 45} |
| Cluster-2 | {1, 8, 41, 9, 23, 28, 29, 31, 42} |
| Cluster-3 | {25, 32} |

### 4.6. Intrinsic protein disorder analysis

In all PAH sequences (Figure 13 (Top)), a prominent presence of highly flexible residues was observed, with an average percentage of 46.74 *±* 3.43. The percentage of disordered residues within each PAH sequence was 13.03, with a standard deviation of 4.54 while the percentage of moderately flexible residues varied between 28.25 and 41.59.

**Figure 13:**
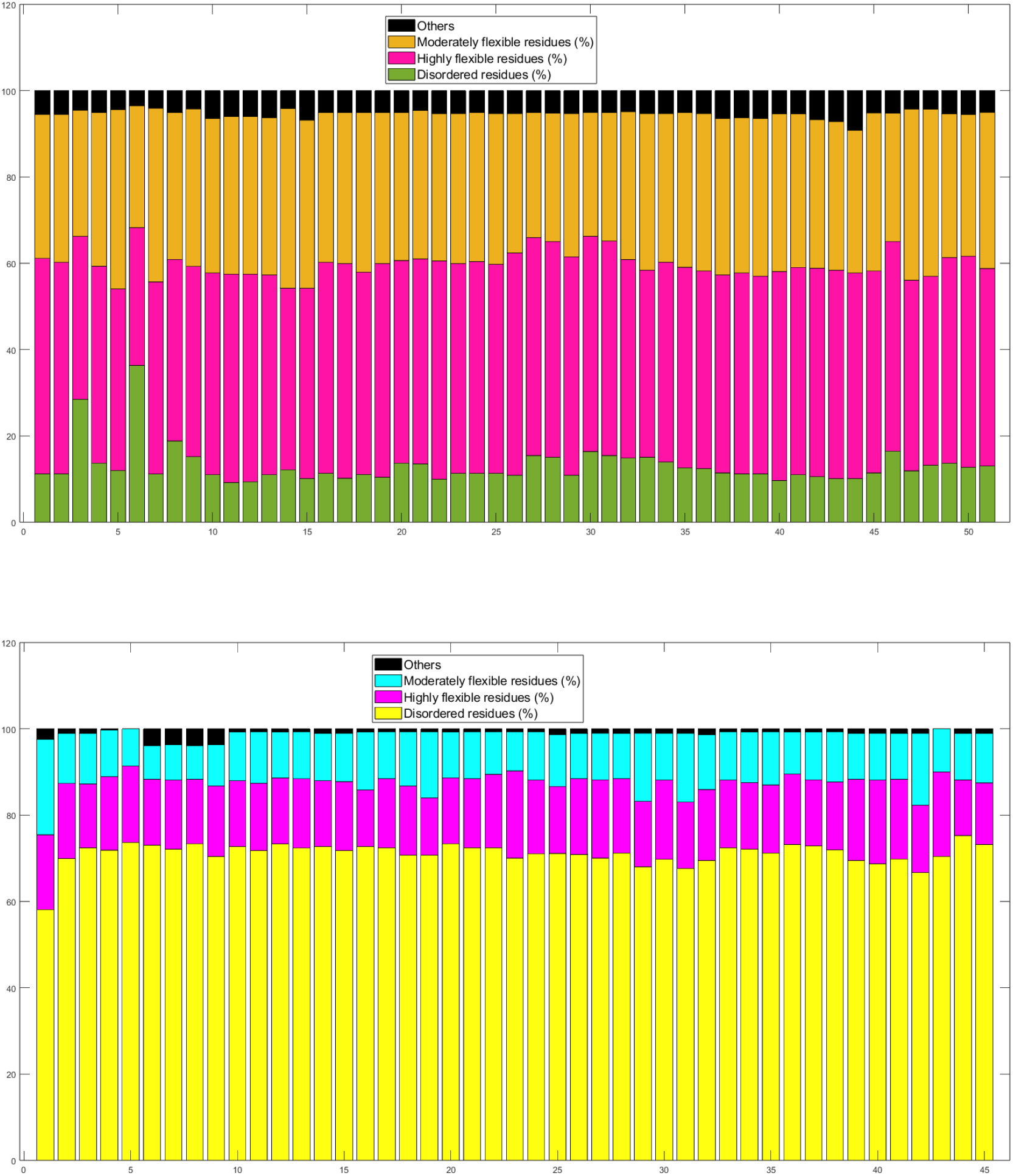
Percentages of disordered, highly flexible, moderately flexible, and other residues in PAH (Top) and SMN1 (Bottom) proteins.

A predominant part of SMN1 sequences (ranging from 58.13% to 71.14%) was identified as the disordered region (Figure 13 (Bottom)). The percentage of highly flexible residues in SMN1 sequences had a mean of 16.24 with a standard deviation of 1.6.

#### 4.6.1. Change response sequences of disordered, highly flexible, moderately flexible, and other residue profiles

In Figure 14 we present the relative frequency distribution of sixteen changes derived from the change response sequence based on the intrinsic disorder profiles of PAH and SMN1 sequences. Notably, we observed that the following transitions did not occur in any of the PAH or SMN1 sequences: O_HF, O_D, MF_D, HF_O, D_O, and D_MF. This absence of transitions between these specific states is of particular significance.

**Figure 14:**
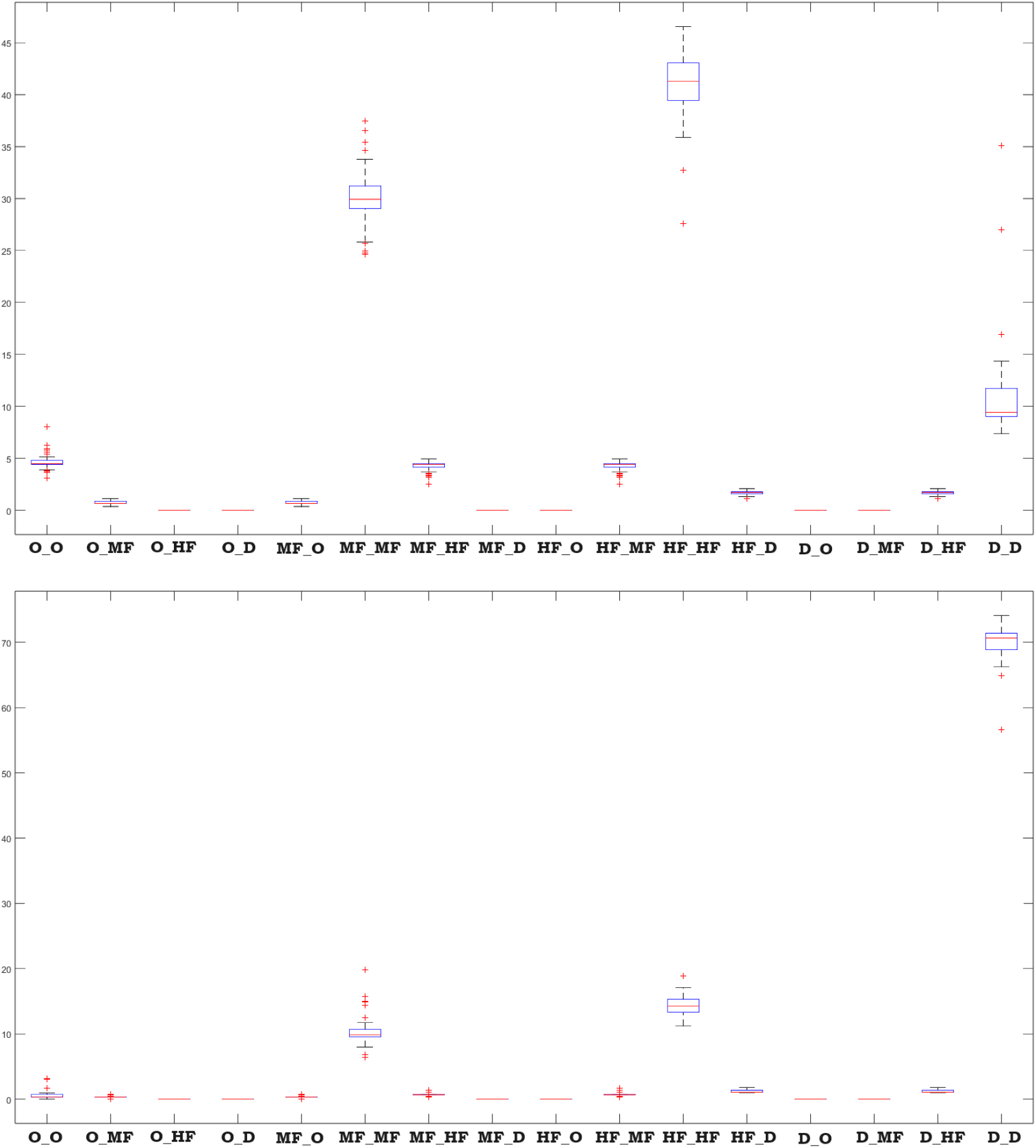
Box-plot of the percentages of disordered, highly flexible, moderately flexible, and other residues changes in PAH (Top) and SMN1 (Bottom) protein sequences.

It is worth mentioning that each PAH protein sequence exhibited the highest proportions of changes from highly flexible to highly flexible residues (HF_HF), with a wide range varying from 27.57% to 46.56%. The next highest percentage of changes involved moderately flexible to moderately flexible (self-transitions) residues in PAH sequences, ranging from 24.63% to 37.47%. This confirms the prevalent presence of consecutive highly flexible residues as well as moderately flexible residues in PAH protein sequences.

Each SMN1 sequence displayed the highest proportions of changes involving disordered to disordered residues (D_D), with a range spanning from 56.59% to 74.12%. Furthermore, it was observed that the percentages of transitions involving moderately flexible (highly flexible) to moderately flexible (highly flexible) residues in SMN1 sequences varied from 6.406 (11.18) to 19.79 (18.919).

Utilizing a distance threshold of 4.5, PAH (SMN1) sequences led to the emergence of four (five) distinct clusters, clearly visible in the dendrogram (Figure 15), summarized in Table 12. Notably, the two most substantial clusters, denoted Cluster-1 and Cluster-2, encompassed 19 and 18 PAH sequences, respectively (Table 12). The largest cluster (Cluster-1) comprised 22 SMN1 sequences (Figure 15 (Bottom)). Here PAH_3 and PAH_6 were outliers among PAH sequences, but SMN_1 was found to be distant from the rest of the SMN1 sequences contrary to previous cases.

**Figure 15:**
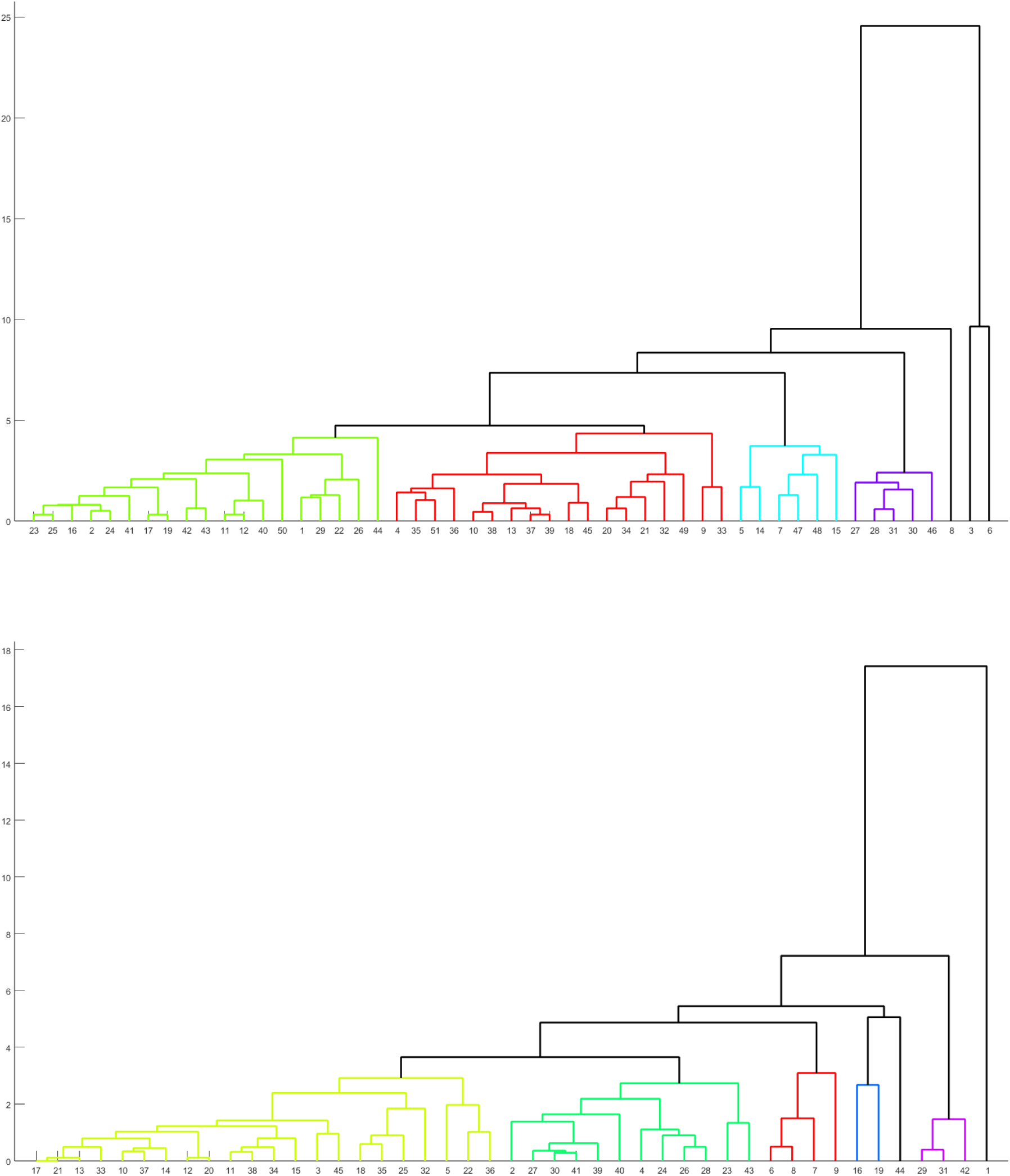
Phylogenetic relationship among the PAH (Top) and SMN1 (Bottom) proteins based on percentages of disordered, highly flexible, moderately flexible, and other residues.

**Table 12:** Clusters of PAH and SMN1 proteins based on the change response profiles of disordered, highly flexible, moderately flexible, and other residues in PAH and SMN1 proteins.

| Clusters | Sequences (PAH) |
| --- | --- |
| Cluster-1 | {23, 25, 16, 2, 24, 41, 17, 19, 42, 43, 11, 12, 40, 50, 1, 29, 22, 26, 44} |
| Cluster-2 | {4, 35, 51, 36, 10, 38, 13, 37, 39, 18, 45, 20, 34, 21, 32, 49, 9, 33} |
| Cluster-3 | {5, 14, 7, 47, 48, 15} |
| Cluster-4 | {27, 28, 31, 30, 46} |
| Clusters | Sequences (SMN1) |
| Cluster-1 | {17, 21, 13, 33, 10, 37, 14, 12, 20, 11, 38, 34, 15, 3, 45, 18, 35, 25, 32, 5, 22, 36} |
| Cluster-2 | {2, 27, 30, 41, 39, 40, 4, 24, 26, 28, 23, 43} |
| Cluster-3 | {6, 8, 7, 9} |
| Cluster-4 | {16, 19} |
| Cluster-5 | {29, 31, 42} |

### 4.7. Phylogenetic relationship based on structural and physicochemical features

#### 4.7.1. I-features

Through a comprehensive analysis using a distance threshold of 7.5 (6), we unveiled the presence of six (three) distinct clusters among PAH (SMN1) sequences based on I-features. The delineation of these clusters is visually depicted in the dendrogram using distinct colors (Figure 16 and concisely summarized in Table 13. Remarkably, for PAH sequences, the largest cluster (Cluster 1) encompassed 22 PAH sequences, and for SMN1, the largest cluster (Cluster 2) contained 16 SMN1 sequences as tabulated in 13. Similar to dendrograms based on the relative frequency of amino acid, PQ profile, and ABN profile, PAH_3 and PAH_6 were distant from most of the PAH sequences while SMN_25, SMN_32, and SMN_43 were away from the rest of SMN1 sequences.

**Figure 16:**
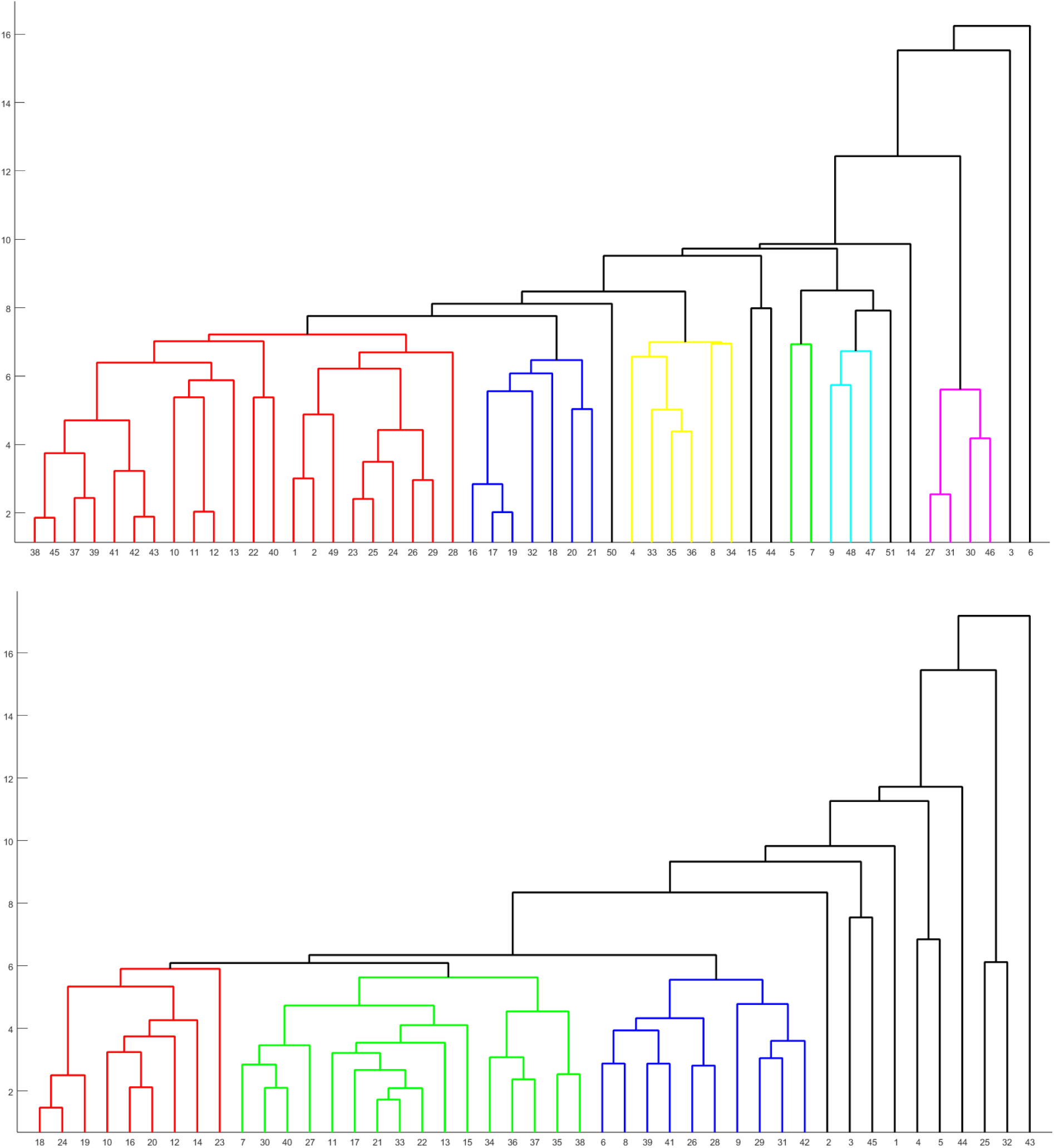
Phylogenetic relationship among the PAH (Top) and SMN1 (Bottom) proteins based on structural and physicochemical features.

**Table 13:** Clusters of PAH and SMN1 proteins based on structural and physicochemical features.

| Clusters | Sequences (PAH) |
| --- | --- |
| Cluster-1 | {38, 45, 37, 39, 41, 42, 43, 10, 11, 12, 13, 22, 40, 1, 2, 49, 23, 25, 24, 26, 29, 28} |
| Cluster-2 | {16, 17, 19, 32, 18, 20, 21} |
| Cluster-3 | {4, 33, 35, 36, 8, 34} |
| Cluster-4 | {5, 7} |
| Cluster-5 | {48, 47} |
| Cluster-6 | {27, 31, 30, 46} |
| Clusters | Sequences (SMN1) |
| Cluster-1 | {18, 24, 19, 10, 16, 20, 12, 14, 23} |
| Cluster-2 | {7, 30, 40, 27, 11, 17, 21, 33, 22, 13, 15, 34, 36, 37, 35, 38} |
| Cluster-3 | {6, 8, 39, 41, 26, 28, 9, 29, 31, 42} |

#### 4.7.2. ProtrWeb-features

Incorporating various ProtrWeb features, as outlined in Section 3.8, the phylogenetic analysis has yielded four clusters comprising a total of 43 PAH sequences based on distance threshold 10 (see Figure 17). It is worth noting that the largest cluster, denoted as Cluster-1, encompasses 28 of the PAH sequences, as indicated in Table 14. Similarly, 37 SMN1 sequences were grouped into six clusters based on distance threshold 8, with the largest cluster, here also Cluster-1, containing 12 SMN1 sequences. PAH_3 and PAH_6 were outliers while SMN_43 were away from the rest of SMN1 sequences.

**Figure 17:**
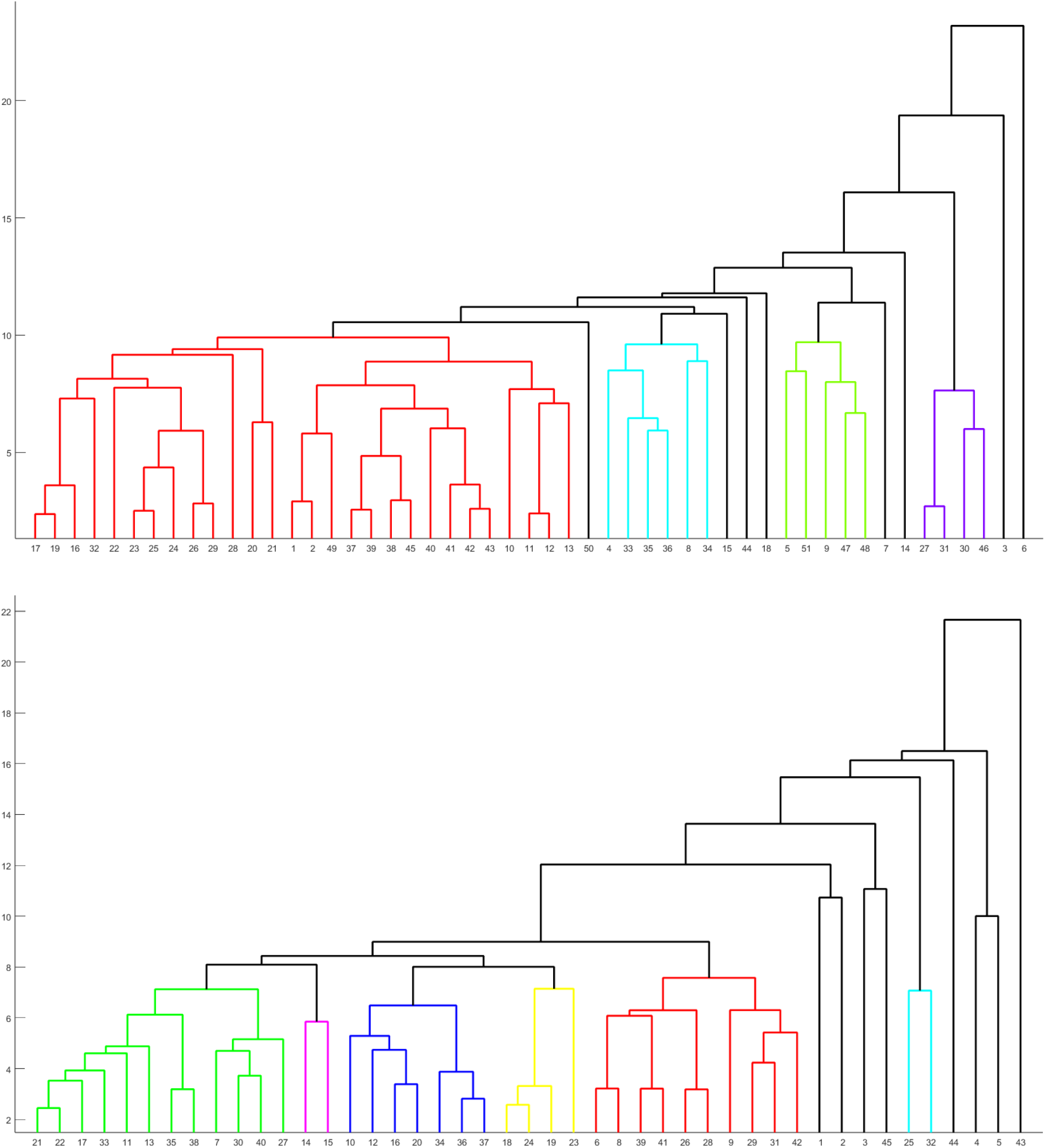
Phylogenetic relationship among the PAH (Top) and SMN1 (Bottom) proteins based on structural and physicochemical features (ProtrWeb).

**Table 14:** Clusters of PAH and SMN1 proteins based on structural and physicochemical features (ProtrWeb)

| Clusters | Sequences (PAH) |
| --- | --- |
| Cluster-1 | {17, 19, 16, 32, 22, 23, 24, 25, 26, 29, 28, 20, 21, 1, 2, 49, 37, 39, 38, 45, 40, 41, 42, 43, 10, 11, 12, 13} |
| Cluster-2 | {4, 33, 35, 36, 8, 34} |
| Cluster-3 | {5, 51, 9, 47, 48} |
| Cluster-4 | {27, 31, 30, 46} |
| Clusters | Sequences (SMN1) |
| Cluster-1 | {21, 22, 17, 33, 11, 13, 35, 38, 7, 30, 40, 27} |
| Cluster-2 | {14, 15} |
| Cluster-3 | {10, 12, 16, 20, 34, 36, 37} |
| Cluster-4 | {18, 24, 19, 23} |
| Cluster-5 | {6, 8, 39, 41, 26, 28, 9, 29, 31, 42} |
| Cluster-6 | {25, 32} |

In summary, structural and physicochemical analyses provide valuable insights into the structural relationships and disparities among the examined PAH and SMN1 sequences. Clustering analyses reveal evolutionary relationships and potential functional similarities, while the identification of structural features informs their roles in biochemical pathways. Variations in physicochemical properties can signify differences in stability, solubility, and interaction capabilities, potentially linking these to unique functions or disease mechanisms in conditions like PKU and SMA. Additionally, these analyses can identify potential drug targets, guiding drug development efforts.

### 4.8. Agglomerated proximal sets of PAH and SMN1 proteins

Proximal sets of PAH and SMN1 proteins were derived as discussed in section 3.10 (Table 15). PAH (SMN1) proteins belonging to the agglomerated proximal sets share a high amount of similarities as extracted from the quantitative features adumbrated in this study.

**Table 15:** List of proximal sets of PAH and SMN1 proteins.

| Serial No. | Proximal Sets (PAH) | Serial No. | Proximal Sets (PAH) | Serial No. | Proximal Sets (SMN1) |
| --- | --- | --- | --- | --- | --- |
| 1 | {1,2} | 7 | {20,32} | 1 | {11,13,17,21,22,33,35,38} |
| 2 | {40,43} | 8 | {33,35} | 2 | {27,30} |
| 3 | {11,12,42} | 9 | {27,30,31,46} | 3 | {10,12,20} |
| 4 | {22,23,24,25,26,29} |  |  | 4 | {34,37} |
| 5 | {13,37,38,39,45} |  |  | 5 | {29,31,42} |
| 6 | {16,17,19} |  |  |  |  |

*Illustration*: Consider *S* = *{*40, 43*}* of PAH proteins. The objective is to show that *S* is an agglomerated proximal set. Note that, *S ⊆ c*_1,2_ (*i* = 1, *p* = 2) (Table 5), *S ⊆ c*_2,2_ (*i* = 2, *p* = 2) (Table 7), *S ⊆ c*_3,2_ (*i* = 3, *p* = 2) (Table 10), *S ⊆ c*_4,2_ (*i* = 4, *p* = 2) (Table 11), *S ⊆ c*_5,1_ (*i* = 5, *p* = 1) (Table 12), *S ⊆ c*_6,1_ (*i* = 6, *p* = 1) (Table 13), and *S ⊆ c*_7,1_ (*i* = 7, *p* = 1) (Table 14). Hence *S* = *{*40, 43*}* is an agglomerated proximal set and consequently, PAH_40 and PAH_43 were derived to be proximal.

## 5. Discussion and Conclusions

Rare diseases like PKU and SMA affect millions of people around the world, effective and affordable treatment strategies are yet to be developed. In the present study, we focus on understanding the correlation between different amino acid changes and the different PAH and SMN1 variants. For both PAH and SMN1, there was a decrease of some order-promoting amino acids and an increase in specific disorder-promoting residues. This correlates with previous studies, in which the SMN1 protein is predicted to have long disordered regions, which are close to the binding sites of SMN1 [80]. Previous studies on PAH have shown that there are also several regions in the PAH sequence, where the disordered regions can also affect the regulatory domain and the protein activity [81]. In our current study, we also found that among all the PAH sequences, the maximum homogeneous polystrings length is 8 and for SMN1 sequences it is 10. Previous studies have shown that the amino acid repeats are crucial for protein function and play an important role in protein-protein interaction [82]. Homogeneous polystring length also provides useful insight into understanding the protein evolution and functional characteristics [83]. Sequences with long polystrings have a faster evolution [83]. Pro-rich polystrings in SMN1, especially those of lengths 8, 9, and 10, characterize the majority of SMN1 sequences, suggesting a potential role of Pro in the structural stability or functional diversity of SMN1 proteins. Additionally, for both PAH and SMN1 the Shannon entropy was very high, suggesting higher degrees of randomness in terms of amino acid frequency, which means that there are more possibilities for the protein conformation [84]. It is important to note that even though Shannon entropy provides a lot of information about the diversity of both SMN1 and PAH, it is also limited in many ways as it does not take into account the environmental and other external interactions [85].

The current study also highlights that both PAH and SMN1 have similar frequencies of polar and non-polar residues, which can be correlated to the importance of stabilizing reaction especially between the polar and non-polar residues [86]. We also investigated the frequency distribution of the PP, NP, PN, NN changes and it was interesting to note that the PAH sequences have a higher percentage of PP change and for SMN1 the highest was the NN residue change. The distribution of the polar and nonpolar residues can affect the protein aggregation and also plays an important role in functional characterisation [60, 87]. Changes in this distribution might affect the protein solubility and function and thereby can lead to disease development.

Both acidic and basic residues play an important role in the functional characterization of the sequence such as DNA interaction, activator, and coactivator interaction, etc. [88]. In the current study, we found that there is a balanced distribution between acidic and basic residues. Such distribution studies are very important as conserved residues are under higher evolutionary pressure and any changes in the acidic and basic amino acid distribution can cause sequence changes and lead to disease phenotypes. Such changes in distribution can also affect the interaction of the SMN1 and PAH with other activators and DNA sequences, thereby leading to misregulation in the pathway. Our study also found that the percentage of NN change of residues was the highest, which can be attributed to the presence of a higher number of neutral residues in both SMN1 and PAH. The phylogenetic relationship derived from spatial distribution of acidic, neutral and basic residues lead to the formation of clusters for PAH and SMN1 sequences which can be correlated with similarity between the different sequences depending on the arrangement of acidic, neutral or basic residues in sequences.

Previous studies have shown that PAH has several flexible residues, is highly flexible in a solution state, and can undergo conformational changes [89, 90]. In this study, we identified an increased presence of highly flexible residues among all the PAH sequences, which also points towards the importance of these flexible residues in conformational changes which can also be mobilized near the active site residues [91]. For SMN1 sequences, there is an increased presence of disordered residues and previous studies have also shown that SMN1 is a highly disordered protein [92]. Identifying such residues becomes very important as changes in the intrinsic protein disorder can cause changes in the conformation, thereby affecting protein activity [93]. This becomes especially important for identifying the disease-causing changes not just at the sequence level but also in spatial conformation. This investigation also highlights the importance of the absence of transition between the different states in SMN1 and PAH sequence for the transitions O_HF, O_D, MF_D, HF_O, D_O, and D_MF. It can be interesting to look further into why such transitions are absent and its potential implications on the protein function and conformation. Studying these signature-genomics can also help us in understanding the interaction of key residues both at sequence level and also spatially.

Furthermore, nine agglomerated proximal sets were identified and cumulatively they include 29 out of 51 PAH sequences, while five agglomerated proximal sets were observed among 18 out of 45 SMN1 sequences (Table 15). Sequences within these proximal sets, whether PAH or SMN1, exhibit a significant level of similarity in terms of the various quantitative features outlined in this study. On the other hand, PAH_3 and PAH_6 were outliers in all dendrograms except dendrograms based on sequence homology where PAH_6 was the only outlier. Among SMN1 sequences SMN_43, SMN_25, and SMN_32 were distant from other SMN1 sequences in four out of seven dendrograms while SMN_43 alone was the outlier in two dendrograms as described in the results section.

This study serves as a pivotal step forward in elucidating the genetic landscapes of SMN1 and PAH, providing a robust foundation for further research aimed at enhancing our ability to comprehend, diagnose, and potentially treat disorders associated with these proteins.

## Author contributions statement

DN, SSH, and ER conceived the problem and theoretical experiments. DN, SSH, VNU, ER, and KL executed the results and performed the analyses. DN, SSH, VNU, KL, and PB wrote the initial draft. All authors reviewed and edited the manuscript. All the authors checked, reviewed, and approved the final version of the manuscript.

## Declaration of competing interest

The authors declare no conflict of interest.

## Notes

### Competing Interest Statement

The authors have declared no competing interest.

